# Mitochondrial carbonic anhydrase-VB inhibition rescues brain endothelial stress and memory in Alzheimer’s disease models

**DOI:** 10.64898/2026.03.16.711716

**Authors:** Nicole L. Lemon, Elisa Canepa, Rafael Vázquez-Torres, Rebecca Parodi-Rullán, Tina Hirt, Lisa M. Petersohn, Tashnuva Rifat, Maryam H. Abyaneh, Marc A. Ilies, Silvia Fossati

## Abstract

Alzheimer’s Disease (AD) is a devastating neurodegenerative disorder with no effective cure, characterized by the cerebral parenchymal and vascular accumulation of aggregated Amyloid-β (Aβ) and hyperphosphorylated tau. Cerebrovascular and mitochondria dysfunction are early causal events in the progression of AD. Previous studies support that inhibiting carbonic anhydrases (CA) may prevent mitochondrial and cerebrovascular dysfunction in AD models. Here, we selectively target the mitochondrial CA isoform CA-VB by pharmacological and genetic manipulation, in human cerebral microvascular endothelial cells (hCMEC) and we confirm the protective effects of the CA-V inhibitor in AD mice. CA-V inhibition and CA-VB KO prevent Aβ induced mitochondria-mediated endothelial apoptosis, loss of barrier resistance, and hCMEC inflammatory activation. Strikingly, CA-V inhibition also mitigates caspase activation and endothelial cell activation in the brains of 3xTg AD mice, resulting in preserved memory function. Our results demonstrate that selective CA-V inhibition is an effective and promising strategy against AD-mediated cerebrovascular pathology, neuroinflammation and cognitive impairment.

## Introduction

Alzheimer’s Disease (AD), the most common form of dementia, is a devastating neurodegenerative disease, currently afflicting over 55 million people worldwide [1]. The most common risk factor for AD and other forms of dementia is age [1].

AD is characterized by the accumulation of extracellular Amyloid-beta (Aβ) aggregates (or plaques) and intraneuronal hyperphosphorylated tau tangles in multiple brain areas, including the hippocampus and cortex [2]. Recently, therapies employing antibodies targeting Aβ to facilitate its clearance have been approved by the FDA for the treatment of AD. However, due to their concerning side effects, such as amyloid related imaging abnormalities (ARIA) [3–6], and only partial efficacy in decreasing cognitive impairment, there is still no effective cure for AD. As the aging population is growing rapidly, it is evident that novel therapies against this devastating disorder are vital.

The demise of the cerebral vessels, with decreased cerebral blood flow (CBF) and increased blood-brain barrier (BBB) permeability, is one of the earliest causal events in the pathogenesis of AD, which in turn affect other cells composing the neurovascular unit (NVU), such as glial and neuronal cells [6–10]. The NVU is a multi-cellular structure composed of cerebral endothelial cells (CEC), pericytes/smooth muscle cells, astrocytes, microglia, and neurons which all work together to execute many vital functions, including maintenance of the physical barrier between the blood and the brain, nutrient exchange, regulation of CBF, and immunological surveillance [2, 11].

Cerebral amyloid angiopathy (CAA), the accumulation of aggregated Aβ around and within the brain vasculature, is observed in about 90% of AD patients [12, 13]. Parallel to NVU dysfunction is the rise of neuroinflammatory processes, which exacerbate a vicious cycle of microvascular stress, BBB permeability, neuroinflammation and glial cell activation, and eventually neurodegeneration [14]. Aβ deposition at the cerebral vessels has been observed to both cause and worsen NVU dysfunction, by affecting mechanisms such as mitochondrial metabolism, oxidative stress, cell death and neuroinflammation [10, 14]. Many studies in mice and humans have confirmed the presence of vascular abnormalities, both Aβ-dependent and accelerating Aβ deposition, very early in the disease [6, 8, 15, 16]. Single-cell transcriptomics in human AD brains has recently established an upregulation of apoptotic pathways in microvascular cerebral endothelial cells [17]. Thus, improving endothelial and cerebrovascular function is a critical target for AD therapies.

In AD brains, Aβ40 is known to accumulate more on the vasculature, while Aβ42 deposits more in parenchymal plaques. Although most cases of CAA are sporadic, there are known familial mutations, such as the Dutch mutation-characterized by Aβ40 mutated in position 22 (AβQ22). The Dutch mutation, among others within the amyloid precursor protein (APP), clinically presents with severe Aβ deposition at the brain vasculature, causing early onset dementia, CAA, strokes, and cerebral hemorrhages [18]. Although wild-type (WT) and Dutch Aβ aggregates induce similar CEC mitochondria-mediated apoptotic mechanisms and BBB abnormalities, as previously shown [19, 20], AβQ22 has a faster aggregation rate, compared to WT-Aβ40 [20–22], correlating with its higher toxicity [19], and therefore it is a useful tool to model CAA toxic effects in cell studies such as this.

Our lab has demonstrated that the FDA approved non-selective, pan-carbonic anhydrase (CA) inhibitors, Acetazolamide (ATZ) and Methazolamide (MTZ) are effective in ameliorating mitochondria-mediated cell death and stress mechanisms in models of amyloidosis [23–26]. In the TgSwDI model of CAA, ATZ and MTZ preserved vascular fitness and prevented gliosis [25]. There is also evidence that CA inhibitors are protective in models of stroke [2], obesity [27], and diabetes [28]. ATZ and MTZ prevent Aβ induced cell death, through both the extrinsic and intrinsic apoptotic pathways, in many cell types, including CEC, smooth muscle cells, glial cells, and neurons, in addition to diminishing mitochondrial oxidative stress [23, 24, 26]. The intrinsic apoptotic pathway, also referred to as mitochondria-mediated apoptosis, is regulated through signaling molecules such as BIM, BAX and BAK, and BCL-2. When BAX and BAK translocate to the outer mitochondrial membrane they form pores, increasing mitochondrial membrane permeability and promoting loss of mitochondrial membrane potential. Mitochondria then release cytochrome C into the cytosol, leading to the activation of caspase-9, which activates caspase-3, resulting in the execution of apoptosis, through DNA fragmentation.

CEC are the physical barrier responsible for the inhibition of entry of peripheral molecules, immune cells, or foreign substances into the healthy brain. When needed, they have a direct role in the recruitment of peripheral immune cells to the brain, as well as in the activation of glial cells within the brain during inflammation. Tight junction proteins (TJPs), such as occludin and claudin-5, are important regulators of BBB integrity. Although there are mixed reports on the expression of TJPs in models of AD, it is clear that the BBB becomes leaky due to AD and CAA pathology. ECs can become activated through the Aβ-mediated accumulation of reactive oxygen species (ROS), or other mechanisms [14]. When EC are activated, they release pro-inflammatory cytokines and increase the expression of immune cell adhesion proteins such as vascular cellular adhesion molecule-1 (VCAM-1), intercellular adhesion molecule-1 (ICAM-1) and E-selectin, which promote the extravasation of immune cells form the periphery into the brain. When this immune response becomes chronic, it is detrimental to the brain tissue [14, 29].

Carbonic Anhydrases (CA) are zinc metalloenzymes that catalyze the hydration of carbon dioxide to produce bicarbonate and a proton. This reaction is vital in many processes, such as the regulation of pH, ion, and redox homeostasis, as well as metabolism [2, 30, 31]. There are 15 different CA isoforms expressed by humans, each one with its’ own tissue expression pattern, activity level, and cellular localization [2, 31, 32], highlighting their different functions. CA isoforms are expressed on the plasma membrane of cells, in the cytosol, and two are expressed in the mitochondria (CA-VA and CA-VB). Importantly, the two mitochondrial isoforms have a differential expression pattern, CA-VB being more abundant in the brain [2, 31, 33].

We have recently observed that the expression of the mitochondrial isoform CA-VB is increased in CEC treated with Aβ, as well as in the brain of the TgSwDI mouse model of CAA, and in human AD and AD+CAA brains, emphasizing the potential role of CA-VB in the development of AD pathology [25], specifically through modulation of cerebrovascular function.

The non-selective CA inhibitors ATZ and MTZ are FDA approved for other indications, such as glaucoma and high-altitude sickness. Although pan-CA inhibitors are demonstrated to be protective i*n vitro* and *in vivo*, chronic long-term use is a concern due to their lack of selectivity and potential undesired effects [2, 34]. ATZ and MTZ belong to the sulfonamide drug class, with established structures which have been modified and screened to determine their inhibitory activity on multiple CA isoforms [32]. In a study evaluating the activity of benzene sulfonamide CA inhibitors, it was determined that the compound 4-phenylacetamidomethyl-benzenesulfonamide (4ITP) has strong selectivity for CA-V [35, 36]. 4ITP is considered highly selective for CA-VA and CA-VB, with a Ki value of 8.6 and 8.3 nM respectively. This is about 10-fold more effective than ATZ and MTZ. 4ITP was determined to be much less effective on cytosolic isoforms CA-I and CA-II, and even less on plasma membrane isoforms CA-IX and CA-XII [35].

Our previous findings demonstrate that pan-CA inhibitors prevent CEC death through mitochondria-specific mechanisms and that CA-VB is increased in AD models and human AD brains. Hence, we hypothesized that the selective inhibition of mitochondrial CA-V would protect from Aβ induced CEC mitochondria-mediated apoptosis and NVU stress, therefore reducing cognitive impairment, while also decreasing the off-target undesired effects of pan-CAIs. Here, we aimed to investigate this hypothesis in CECs as well as the 3xTG mouse model of AD [37], which presents both Aβ and tau pathology, in addition to vascular dysfunction [38].

This study establishes that a known selective CA-V inhibitor, 4ITP, is effective at reducing Aβ-induced mitochondria-mediated CEC death, BBB dysfunction and cerebrovascular stress *in vitro*. We also confirmed the specific role of CA-VB in Aβ-mediated CEC death and BBB dysfunction, through CRISPR Cas9 CA-VB knockout (KO) in human CEC. Moreover, we pioneered the employment of this CA-V inhibitor in an *in vivo* AD model, showing that it can prevent endothelial cell loss and BBB dysfunction, mitigate neuroinflammation, and ameliorate cognitive impairment in 3xTG mice, therefore shedding light on CA-V as a potential specific target for AD therapy.

## Materials and Methods

### Cell Culture

Immortalized human cerebral microvascular endothelial cells (hCMEC/d3) were provided by Babette Weksler (Cornell University). Cells were grown and maintained in endothelial basal medium (EBM-2) (Lonza CC-3156) supplemented with EGM-2 growth factors (Lonza CC-4176) and 5% fetal bovine serum (FBS).

CRISPR Cas9 mediated KO cell pool of CA-VB in hCMEC/D3 cells were generated by EditCo Bio, Inc. (Redwood City, CA, USA). To generate these cells, Ribonucleoproteins containing the Cas9 protein and synthetic chemically modified guide RNA produced by Synthego were electroporated into the cells using EditCo’s optimized protocol. Editing efficiency is assessed upon recovery, 48 hours post electroporation. Genomic DNA is extracted from a portion of the cells, PCR amplified and sequenced using Sanger sequencing. The resulting chromatograms are processed using EditCo’s Inference of CRISPR edits software. The wild-type cell line was generated similarly without gRNA specific to the gene of interest.

Upon receiving the cells and prior to experiments, CA-VB KO was confirmed via protein expression. Cells were grown and maintained in EBM-2, supplemented with EGM-2 growth factors and 5% FBS.

Cells were grown in a humidified cell culture incubator, under a 5% CO_2_ atmosphere at 37°C. WT hCMEC were routinely checked to ensure that they maintained their proper function and TJP expression, as CRISPR/Cas9 CA-VB KO to ensure CA-VB KO stability.

### Synthesis and characterization of the CA V selective inhibitor 4ITP

The following materials were used as received: phenylacetic acid, thionyl chloride were from Sigma-Aldrich (St. Louis, MO), 4-aminomethylbenzenesulfonamide hydrochloride was from Oakwood Chemical (Estill, SC), N-methylmorpholine was from TCI America (Portland, OR). Organic solvents (HPLC quality) were purchased from Fisher Scientific (Pittsburgh, PA), EMD (Gibbstown, NJ), and VWR International (West Chester, PA).

The purity and the structure identity of the intermediary and final products were assessed by thin-layer chromatography (TLC), HPLC-MS, ^1^H-, ^13^C-NMR. TLC was carried out on SiO_2_-precoated aluminum plates (Alugram® SIL G/UV_254_ 20×20 cm with F254 indicator; layer thickness 0.20 nm; pore size 60 Å) from MACHEREY-NAGEL, (Bethlehem, PA). Flash chromatography was performed using an Isco Combiflash RF (Teledyne Isco, Lincoln, NE), using Isco prepacked silica columns. The purity of compounds was also assessed via LC-MS, using an Agilent 1200 HPLC DAD-MS system (Santa Clara, CA) equipped with a G1315A DAD and a 6130 Quadrupole MS via a ZORBAX SB-C18 column eluted with H_2_O (0.1% HCOOH)/MeCN (0.1% HCOOH) 95/5 to 0/100 linear gradient. NMR spectra were recorded at T = 300 K with a Bruker Avance III 400 Plus spectrometer equipped with a 5 mm indirect detection probe, operating at 400 MHz for ^1^H NMR, at 100 MHz for ^13^C NMR. Chemical shifts are reported as δ values, using tetramethylsilane (TMS) as internal standard for proton spectra and the solvent resonance for carbon.

4-(Aminomethyl) benzenesulfonamide hydrochloride (1 eq, 4.5 mmol, 1.00 g) was suspended in dry acetonitrile (18 mL) in a 50 mL round bottom flask and the flask was purged with argon. The mixture was cooled to 0°C using an ice bath, then 4-methylmorpholine (2.1 eq, 9.4 mmol, 0.95 g) was added under stirring. After 30 min, the ice bath was removed, and the reaction mixture was stirred overnight at room temperature. Separately, phenylacetic acid (1.05 eq, 4.73 mmol, 0.73 g) was treated with thionyl chloride (10 eq, 45 mmol, 5.35 g) and 1 drop of dimethylformamide in a 100 mL round bottom flask and the mixture was heated to reflux, under stirring, for 1h. Excess thionyl chloride was removed in vacuo using a rotary evaporator and the acid chloride was dissolved in dry acetonitrile (3 mL) and added dropwise to the previously made solution of 4-(aminomethyl)benzenesulfonamide, at 0°C (ice bath cooling). The reaction was stirred at 0°C for 30 min, then at room temperature overnight, when a precipitate was formed. The suspension was filtered and the precipitate was washed subsequently with acetone and water, dried under vacuum, and absorbed onto SiO_2_ using MeOH/CHCl_3_. The product was purified by flash chromatography on silica gel (0-40% MeOH/CHCl_3_ gradients). The pure fractions were combined and evaporated to dryness to yield a while solid (55-71% (multiple batches made).

4ITP: ^1^H-NMR (400 MHz, DMSO-d^6^, δ, ppm): 8.63 (t, *J* = 5.8 Hz, 1H, NH), 7.74 (d, *J* = 8.4 Hz, 2H, H_2,6_-PhSO_2_NH_2_), 7.3 (d, *J* = 8.4 Hz, 2H, H_3,5_-PhSO_2_NH_2_), 7.20-7.35 (m, 5H, PhCH_2_CO), 7.29 (s, 2H, SO_2_NH_2_), 4.32 (d, *J* = 5.9 Hz, 2H, NHCH_2_), 3.49 (s, 2H, CH_2_CO). ^13^C-NMR (100.6 MHz, DMSO-d^6^, δ, ppm): 170.2, 143.5, 142.5, 136.2, 128.9 (2C), 128.1 (2C), 127.3 (2C), 126.3, 125.5, 42.2, 41.7.

4ITP: LC-MS (ESI) 305.1 (MH^+^), t_R_= 2.47 min, > 96% purity).

### Determination of 4ITP solubility in water and PBS buffer

The solubility of 4ITP in water and in PBS buffer was determined by the same HPLC method and instrument described above. A stock solution of 4ITP at a concentration of 10 mM were prepared in DMSO and serial dilutions (5 mM, 2.5 mM, 1.25 mM, 0.625 mM, 0.3125 mM and 0.156 mM) were done with the same solvent. A volume of 1 µL of these 4ITP solutions was injected in the instrument and the resulting peak areas were plotted against the corresponding concentrations to obtain a calibration curve, via the least squares linear regression analysis. 4ITP saturated solutions in deionized water and in PBS buffer were made by suspending 10 mg pure 4ITP in 1 mL water or in 1 mL PBS at pH 7.4 in two Eppendorf tubes, vortexing the suspensions for 1 min and subsequently placing them in a mini rotator (Labnet International, Edison, NJ) overnight. The next day the undissolved compound was centrifuged down using a bench centrifuge for 5 min and 10 µL of each supernatant was collected in an LCMS vial. A volume of 1 µL of these 4ITP solutions was injected in the HPLC instrument and the concentration of 4ITP in each solution was determined from the resulting peak areas via the previously constructed calibration curve. The analysis was done in triplicate and the values were reported as mean ± standard deviation.

4ITP solubility in water: 0.057 ± 0.011 mg/mL (0.187 mM) 4ITP solubility in PBS (pH = 7.4): 0.0399 ± 0.003 mg/mL (0.13 mM)

### 4ITP lipophilicity (Log P) determination

Log P value of 4ITP was determined using the same HPLC method and instrument described above, using an injection volume of 1 µL and a flow rate of 1mL/min in all cases. DMSO solutions (1 mg/mL) of reference compounds of known log P (indomethacin, caffeine, ketoprofen, phenytoin, testosterone) were injected into the HPLC system and the retention time of each compound was determined in the same RP C18 column. A calibration curve was constructed by plotting the log P values of these reference compounds against their retention time [39]. Subsequently, 1 mg 4ITP was dissolved in 1 mL of DMSO and was injected in the same system to determine the retention time, which was interpolated within the calibration curve to determine the log P value of 4ITP.

4ITP Log P Value: 1.69

### 4ITP solubilization and delivery in vitro

CA-V inhibitor, 4ITP, was dissolved in DMSO at a concentration of 10mM and stored at-20°C. On the day of treatment, the 10mM stock concentration was diluted to 2mM in EBM2 with 50%FBS. Finally, 4ITP was diluted to its final concentration (1, 10, or 30μM) in treatment medium, EBM2 with 1% FBS.

### Aβ preparation

Aβ40-Q22 was synthesized by peptide 2.0. To obtain monomeric Aβ40-Q22, the peptide was weighed and dissolved in 1,1,1,3,3,3-169 hexafluoro-2-propanol (HFIP; Sigma, St. Louis, MI, USA), at a concentration of 1 mM, and incubated overnight at room temperature to dissolve beta-sheet structures. The day after, the peptide was flash-frozen in dry ice-cold 100% ethanol, then lyophilized until the pellet was dry, and stored at-20°C. To maintain the peptide in its monomeric form, the pellet was resuspended at 10mM in DMSO and then diluted to 1mM with sterile dH_2_O immediately before treatment. Aβ40-Q22 was brought to a final concentration of either 25 or 10µM in EBM2 with 1% FBS for treatment and added to the cells.

### DNA fragmentation/Apoptosis

Cells were seeded at a density of 20,000 cells/well in a 24-well plate 24 hours (h) prior to treatment in complete cell culture media. Following treatments, cells were centrifuged at 200xg for 10 minutes, and the appropriate lysis buffer was then added (Roche Applied Science). DNA fragmentation was measured using the Cell Death ELISA kit (Roche Applied Science), as recommended by the manufacturer. Absorbance was measured with SpectraMax I3x Multimode micro-plate reader (Molecular Devices) at wavelength 405 nm. Absorbance values were analyzed by determining the relative change to the control through the calculation of fold change.

### Cell Viability Assay

Cells were seeded at a density of 10,000 cells/well in a 96-well plate in complete cell culture media 24h prior to treatment. Cells were treated in EBM-2 supplemented with 1% FBS plus the desired concentrations of Aβ40-Q22 and 4ITP. Cell viability was measured after 24h of treatment. WST-1 reagent (Roche, Applied Bioscience) was added directly to the cell culture media (1:10 ratio), and then incubated for 3 hours at 37°C to allow for the colorimetric reaction. Absorbance was read with SpectraMax I3x at a wavelength of 450 nm.

### Cell Event Caspase 3/7

A day before treatment, 10,000 cells/well were seeded in a 96-well plate in complete cell culture media. Cells were treated in EBM-2 supplemented with 1% FBS and the experimental concentrations of Aβ40-Q22 and 4ITP. After 24h of treatment, the Cell Event Caspase 3/7 reagent (Thermo Fisher Scientific) was diluted in Hank’s balanced Salt Solution (HBSS) at a concentration of 5μM, added at a volume of 100μl/well to the cells and incubated at 37°C for 30 minutes. The reagent was then removed and replaced with HBSS during acquisition. Pictures were acquired on an EVOS M5000 microscope at 10x magnification. Images were acquired with the brightfield and GFP channels to visualize the total number of cells and the caspase 3/7 activity, respectively. Three pictures per well were taken and then analyzed using Fiji/ImageJ software. The number of active caspase 3/7 positive cells was normalized to the total number of cells in each image and expressed as percentage of caspase 3/7 positive cells.

### Western blot

In hCMEC, one day prior to treatment, cells were counted and seeded in a 6-well plates at a density of 350,000 cells/well. Following treatment, proteins were isolated from cells. Briefly, media was collected, and cells were washed with 1X PBS and lysed with RIPA buffer (Invitrogen) + Halt protease/phosphatase inhibitors (Thermo Fisher Scientific). Cell debris were pelleted at 16,000xg for 20 minutes at 4°C. Supernatant was collected, and protein concentration was quantified with BCA method, and normalized to 25µg in each sample.

For WB analysis, cell lysates, and homogenates from hippocampus and cortex of WT, 3xTG and 3xTG treated with 4ITP were prepared using BOLT 4x loading buffer and 10x reducing agent or Laemmli Buffer 6x (Invitrogen), depending on which gel was used. To further denature the samples, they were incubated for 5 minutes at 95°C.

Proteins were separated using 4-12% BOLT bis-tris SDS polyacrylamide gels (Invitrogen) or 10% CRITERION SDS polyacrylamide gels (BioRad) and transferred onto 0.45 µm nitrocellulose membrane using 1x Towbin buffer containing 20% methanol, at 110 volts for 70 minutes. After transfer, membranes were blocked with 5% milk in TBS. Primary antibodies, Caspase-9 (1:500 ab202068), Bax (1:500 Novus Biologicals NBP1-88682), Bcl-2 (1:500 ab196495), Bim (1:500 ab32158), Bak (1:250 ab104124), CA-VB (Novus Biologicals NBP1-86090), Occludin (1:250 Invitrogen 71-1500), Claudin-5 (1:500 Invitrogen 352500), ICAM-1 (1:500 Invitrogen MA5407), VCAM-1 (1:500 ab134047), and actin (1:3000 Millipore MAB1501) were incubated on the membrane O/N at 4°C or for 2h room temperature. Membranes were then washed with TBS-T (0.1% Tween) and incubated with the species-appropriate secondary antibody (Licor, 1:20,000 in TBS) for 1h at room temperature. Images were acquired using LICOR Odyssey CLx Immunoblot imager and then analyzed with LICOR Image Studio Software. Uncropped membranes of each representative WB have been provided in a supplementary file.

### Cytochrome C and Mito-tracker Staining

Prior to seeding, 8-well chamber glass slides were pre-coated with attachment factor (Cell Systems). For immunocytochemistry experiments, 15,000cells/well were seeded 24h prior to treatment. Cells were treated with 25µm Aβ40-Q22 with or without 4ITP. After 16h of treatment, Mito-tracker CMH2xROS (Invitrogen) was diluted in 1% FBS EBM2 (1:2000) and added to the cells for 30 minutes at 37°C. This is a dye that only enters mitochondria with a healthy membrane potential. After live-cell staining was complete, cells were washed 1x with PBS and fixed with 4% PFA for 15 minutes at 37°C, then washed again 3x with PBS for 5 minutes. Cells were then permeabilized with 0.2% Triton in PBS for 10min at room temperature, followed by one hour of blocking with 3% BSA in PBS. Then, cells were incubated with CytochromeC-AlexaFlour488 antibody (BD Pharmigen 560263) for 1h at room temperature. Finally, to visualize the nucleus, we used DAPI mounting media (Southern-Biotech 0100-20). Images were acquired using a Nikon Ti2-E fluorescence deconvolution microscope at 100x resolution.

### Mitochondrial H_2_O_2_ production

HCMEC were seeded at 2 million cells/well in 10cm dishes 24h prior to treatment and then treated with Aβ40-Q22 with or without 4ITP. After 16h of treatment, mitochondria were isolated in an isotonic buffer including 75 mM sucrose, 10 mM Tris-HCl pH = 7.4, 2 mM EGTA and 225 mM Mannitol (STEM) buffer [24]. The mitochondrial fraction was quantified using BCA assay. H_2_O_2_ was measured with an Amplex red assay (Invitrogen). Levels of H_2_O_2_ were measured by absorbance detected at 560 nm, normalized to protein quantification of the mitochondria fraction, and then expressed as fold of change to the control.

### Lipid peroxidation 4-HNE ELISA

The levels of 4-Hydroxynonenal (4-HNE) in total cell lysates were measured with a commercially available ELISA kit (Abcam 238538) as per the manufacturer’s instructions. Protein samples were collected after 16h of treatment and then flash frozen with dry ice. The levels of 4-HNE were calculated through a standard curve, detected with Spectra I3x Max at a wavelength 450 nm. Concentrations were further converted to %change of control.

Mouse cortices were homogenized following brain tissue dissections. The same commercially available kit (Abcam 238538) was used for samples *in vivo* normalized to equal mg of protein.

### Measurement of barrier resistance by ECIS Z*𝜽𝜽*

Trans-endothelial electrical resistance was measured using the ECIS ZΘ system (Applied Biophysics). All experiments were performed on 8-well ECIS (8WE10+, Applied Biophysics) 40-electrodes-gold plated arrays, pre-coated per the manufacturer’s instructions. Cells were seeded at a density of 200,000 cells/well, and given 48h to form a monolayer, visualized on the recorded plot as a plateau in Resistance over time. The frequency applied to the cells was 4000 Hz. At this point, cells were treated with 10µM Aβ40-Q22, with or without 4ITP, or 4ITP alone. The resistance was measured for 48h after treatment. Data were expressed as normalized to the untreated control, over a 48-hour period.

### Meso-Scale Discovery Cytokine analysis

Cytokine release was evaluated in conditioned media from untreated hCMEC, and Q22-treated cells with or without 4ITP. Conditioned media (900µl) was collected and then centrifuged at 800 x g for 5 minutes at 4°C. Conditioned media was then flash frozen and stored at-80 for no longer than 1 month. MesoScaleDiscovery (MSD) V-PLEX proinflammatory cytokine panel 1 was used to evaluate 10 cytokines (IL-1β, IL-2, IL-4, IL-6, IL-8, IL-10, IL-12p70, IL-13, TNFα, and IFN-γ). The technique employs an electrochemical detection method which produces a specific signal in order to evaluate all 10 cytokines in one sample/well. The amount of cytokine was calculated using a standard curve, normalized to the amount of protein in the cell sample, and then further converted to FOC of the untreated control.

We also analyzed cytokines and chemokines *in vivo*. In mouse brain cortex homogenized in tissue homogenization buffer (THB) we normalized samples to equal protein amounts of 140ug/well. The samples were loaded, and the protocol was followed per manufacturer’s instructions.

### qPCR array to measure cytokine mRNA levels

HCMEC were seeded 1-day prior to treatment in 6-well plates at a density of 350,000 cells/well. Cells were treated for 24 hours with amyloid, plus or minus CA-V inhibitor 4ITP. Conditioned media was removed and then the cells were washed with 1x PBS. Next, total RNA was extracted using miRNeasy mini kit (Qiagen). Following total RNA purification, samples were quantified with nanodrop and normalized to 1ug/sample. This was followed by a 20µl reverse transcription reaction using Super Script IV VILO master mix to obtain cDNA. The cDNA samples were further diluted with 150ul of nuclease-free water. Next, samples were prepared with taq-man fast advanced master mix and loaded onto the customized Taq-man qPCR array. The targets analyzed were CCL2, CCL3, CXCL1, CXCL2, CSF-2/GM-CSF, CSF3, IL-10, IL-6, IL-12a, IL-12b, IL-4, VCAM-1, SELP/P-selectin, SELE/E-selectin. Results were analyzed by normalizing the target genes to cyclophilin-D/PPIB, then further calculating the FOC of the non-treated control.

### Mouse model and treatment

All procedures and experiments on animals were conducted and approved in accordance with Temple University Animal Care and Use Committee (IACUC) guidelines. The 3xTG AD mouse model was used to evaluate the protective effects of CA-V inhibition *in vivo*. The 3xTG mouse model carries human Amyloid Precursor Protein (APP) Swedish, Presenilin 1 (PSEN1 M146V) and tau (MAPT P301L) transgenes under the Thy1.2 neuronal promoter, and thus presents both Aβ and tau accumulation within the brain, particularly in the hippocampus and cortex, with reports of Aβ accumulation within the vasculature [40]. Mice are anticipated to present with mild amyloidosis around 6 months of age, while tau accumulation is reported later, around 12 months of age [37]. Recent studies have demonstrated that vascular abnormalities [38], metabolic deficits, and chronic inflammation may occur in these mice prior to protein accumulation, particularly more in females than males, translating more accurately to the human progression of disease [38, 41]. In this study, we compared male and female WT mice (C7BL/6;129X1/SvJ;129S1/Sv) (N=20), WT + 4ITP treated mice (N=11) 3xTG mice (N=21), and 3xTG + 4ITP (N=20) treated mice. 4ITP treatment in 3xTG and WT mice began at 6 months of age until 16 months. CA-V inhibitor, 4ITP, was provided in the mice’ diet at 20mg/kg/day, (100 ppm) in a grain-based diet. This was done in collaboration with bio-Serv. Food was stored in a cool dry place. Body weight and activity levels were monitored and recorded throughout the study.

### Barnes Maze

When mice reached 15 months of age, spatial memory was evaluated through the Barnes Maze. The Barnes maze runs over the course of 7 days: day 0, habituation, days 1-5 training and day 6 probe day. On habituation day, mice were put into the middle of the maze with a beaker, a white noise at 80 db for 60 seconds, after which, mice were guided to the escape hole box and left there for 1 minute with the noise off. After habituation day (0), every trial day (1-5), mice underwent 2 trials, 2h apart from each other, in which the mouse had 3 minutes to find the escape hole box. The number of mistakes, in addition to the time to find the escape hole (latency) was recorded on each trial. Between mice, the maze was cleaned, and each day the platform was rotated. On probe day (6), the mice are put into the maze, with the escape box removed. Number of mistakes and the time to find escape hole (when their nose is placed into the correct hole) was recorded with ANY-maze software.

### Fear Conditioning

To assess cue-induced memory, we employed an optimized cue-induced fear conditioning protocol. On Day 0 (habituation), mice were introduced to the conditioning chamber for 5 minutes without any stimulus. The following day (acquisition/training), mice were placed in the same chamber and exposed to a series of tone-foot shock pairings. The foot-shock was set to 0.5mA/2 seconds. Following training, mice returned to their home cage. On day 2 (cue-induced testing phase) mice were placed back into the same chamber and exposed to the same tone without foot shocks. The freezing time in both the acquisition phase (day1) and cue-induced testing phase (day2) were recorded in order to assess cue-induced memory recall. Freezing time was defined as the absence of all movements other than respiration and was recorded with ANY-maze software to avoid human bias.

### Novel Object Recognition Test

Recognition memory was evaluated using a novel object recognition paradigm. On day 0 (habituation) mice were allowed to explore the empty arena for 5 minutes. On day 1, Trial 1 (training phase) mice were placed into the chamber with two identical objects (towers) and allowed to explore for 10 minutes. Two hours later, on day 1, trial 2 (testing phase) one of the familiar objects was replaced with a novel (dice) and placed into the chamber. Mice were allowed to explore for 3 minutes during the testing phase. The location of the novel object was alternated throughout the study to ensure there was no biased in cage side.

All sessions were video-recorded and the time spent focused on each object was captured through ANY-maze software. The %time exploring the novel object during the testing phase was calculated as (time spent exploring the novel object / total exploration time) *100. Any mouse that did not explore the objects for more than 10 seconds was excluded from analysis.

### Rotarod

Locomotor function, coordination and balance were evaluated using an accelerating rotarod procedure across six trials spaced by 20-minute inter-trial interval (ITI). Five mice were run simultaneously on 9.5 cm diameter rods of a land rotarod apparatus. All mice were first given a habituation trial where they were required to stay on the platform for 1 minute without falling. Animals were then tested in experimental trials, during which the rod was accelerating steadily from 4 to 40 rpm over the course of 5 minutes. The latency to fall from the platform was recorded.

### Open-field test

Locomotor activity and exploration were evaluated through open-field test. This test evaluates the distance travelled by the mice, as well as the preferred behavior of mice to spend more time exploring around the perimeter of the box rather than in the center. More time spent in the center suggests decreased anxiety. To start, mice were placed in the center of a rectangular box and then recorded for one-30-minute session. After 30 minutes of exploration mice returned to their home cage. The percentage of time spent in the center as well as the total distanced traveled was recorded.

### Tissue collection and mouse brain processing

Following behavioral analysis, mice were anesthetized and perfused with ice-cold phosphate-buffered saline (PBS). Each brain was split into hemispheres; one hemisphere was fixed with 4% paraformaldehyde and then further processed for immunohistochemistry analysis. The other hemisphere was dissected into pathological brain areas such as the cortex and hippocampus. After dissection, tissue was flash frozen and stored in-80°C.

The tissue for immunohistochemistry was fixed for 2 days in 4% paraformaldehyde at 4°C. Following this, brains were placed in 15% sucrose for 1 day and then 30% sucrose for two. Following fixation and sucrose, brains were washed with PBS and then assembled in a plastic mold with Tissue-Tek O.C.T compound (Fisher Scientific). Brains were stored until sectioning at-80°C.

For WB, hippocampus, and cortex tissue (WT, 3xTG, 3xTG + 4ITP N=6/group males and females) was homogenized in Tissue Homogenization Buffer (20mM Tris-Base, 0.25M sucrose, 5mM EDTA, 1mM EGTA + HALT protease/phosphatase inhibitors). Protein content of the homogenates was quantified with BCA and then normalized to 30μg of protein. WB samples were prepared with 6x Laemmli Buffer (Invitrogen).

### Extraction and quantification of soluble/insoluble amyloid and tau

Hippocampal and cortical brain samples were homogenized in THB buffer (as previously explained) and then used to fraction the soluble and insoluble proteins in 3xTG treated and non-treated mice as previously published [25]. After homogenization, samples were quantified and normalized to equal levels of protein. To obtain soluble protein containing Aβ and tau monomers, oligomers, and protofibrils a diethylamine (DEA) extraction was performed. Additionally, formic acid was employed to obtain the insoluble fraction containing aggregated proteins such as Aβ and hyperphosphorylated tau. Both extracted fractions were then used in specific ELISA (Invitrogen) assays to measure the amount Aβ40, Aβ42, and pTau231.

### Immunohistochemistry

Brains were sectioned at 8-and 20-micron thicknesses with a cryostat. Brain sections were collected on positively charged microscope slides and stored at-80°C until used for staining. For immunostaining evaluation, brain sections were fixed in 4% paraformaldehyde for 20 minutes at room temperature. Next, brain sections were blocked with 10% NGS and 1% BSA solution for 2 hours at room temperature. Then, sections were incubated overnight at 4°C with primary antibodies diluted in 0.3% Triton X-100 (Sigma) blocking solution. The following primary antibodies were used: chicken anti-GFAP (1:1000 Aves #GFAP), rat anti-CD31 (1:100 BD Pharmingen 55370), rabbit anti-Albumin (1:500 Invitrogen PA5-85166), conjugated Alexa fluor 647-cleaved caspase-3 (1:100 Bioss bs.20363R-bf647) and conjugated Alexa fluor 488-NeuN (1:100 Millipore ABN78A4). After overnight incubation, tissue was washed with 1x PBS and then the species-appropriate Alexa-fluor conjugated secondary antibody diluted 1:1000 in PBS were placed on the tissue for 2 hours at room temperature. Further, nuclear staining with DAPI (Invitrogen 1.5ug/ml) was placed on the tissue for 10 minutes at room temperature. Next, sections were washed with PBS and finally coverslips were placed with mounting media.

Stained sections were imaged with a Nikon Ti2-E fluorescence deconvolution 860 microscope equipped with 340/380, 465/495, 540/580 and 590/650 nm excitation filters, keeping identical settings within each session, and using either 10x, 20x, or 60x zoom objectives. 20x and 60x images were acquired with a 0.5μm Z-stack. To have a consistent examination for each animal, 5-10 different images were acquired in the same brain area of interest. Images were deconvolved and then Z-stack images were merged into 2D maximum signal projections of equal thickness using Fiji. All image analysis was conducted using the object colocalization FL module in HALO Image Analysis Platform version 4.2.6399 (Indica Labs, Inc).

### Vessel diameter measurement

Vessels were stained using CD31 antibody described above using 20-micron thick tissue sections. After image acquisition and image processing, 3 images per mouse, per brain area (hippocampus and cortex) were selected. 20 vessels/image were outlined via the annotation tool within HALO software. We then obtained 6 diameter measurements/vessel. These 6 values were averaged to obtain each blood vessels avg. diameter. Each vessel’s diameter was graphed allowing us to visualize the shift in the population of blood vessels within each group. Only measurements for micro-vessels (<10 microns in diameter) were analyzed.

## Results

### CA-V inhibition reduces mitochondria-mediated Aβ-induced apoptosis in cerebral endothelial cells

To confirm that selective inhibition of mitochondrial CA-V prevents CEC death and mitochondria-mediated apoptotic mechanisms [23–26, 34], we treated hCMEC with 25µM AβQ22 alone or in combination with the CA-V inhibitor 4ITP (1,10,30µM) for 24h **(Figure 1)** or 16h **(Figure 2)**. These time points were selected based on previous evidence demonstrating AβQ22’s effects on mitochondria-mediated cell death within hCMEC [24]. After confirming that AβQ22 induced apoptosis, measured as DNA fragmentation (the last step of the apoptotic pathway), we determined that 10 and 30µM 4ITP significantly protected hCMEC from AβQ22-induced apoptosis **(Fig 1A)**, nearly rescuing DNA fragmentation, as well as CEC loss of viability (**Fig 1B),** to the control levels. Since both concentrations (10 and 30µM) of 4ITP were non-toxic to hCMEC **(Suppl. Fig 1)** and rescued both AβQ22-induced hCMEC DNA fragmentation **(Fig 1A**) and viability loss **(Fig 1B)**, these concentrations were selected for subsequent experiments. To test whether CA-V inhibition could prevent Aβ-induced mitochondria-mediated apoptosis, we measured AβQ22-induced caspase-9 activation, which is directly promoted by mitochondrial Cytochrome C release. We demonstrated that 10 and 30µM 4ITP were protective against AβQ22-induced caspase-9 activation **(Fig 1C)**, assessed as increased formation of its cleaved fragment by WB. CA-V inhibition also significantly reduced the percentage of active executioner caspases 3 and 7-positive cells **(Fig 1D),** measured by the specific fluorescent marker Cell-Event caspase 3/7 (Invitrogen). These results suggest that selective CA-V inhibition with 10 and 30µM 4ITP has a protective effect against AβQ22-induced mitochondria-mediated CEC apoptosis.

**Figure 1.**
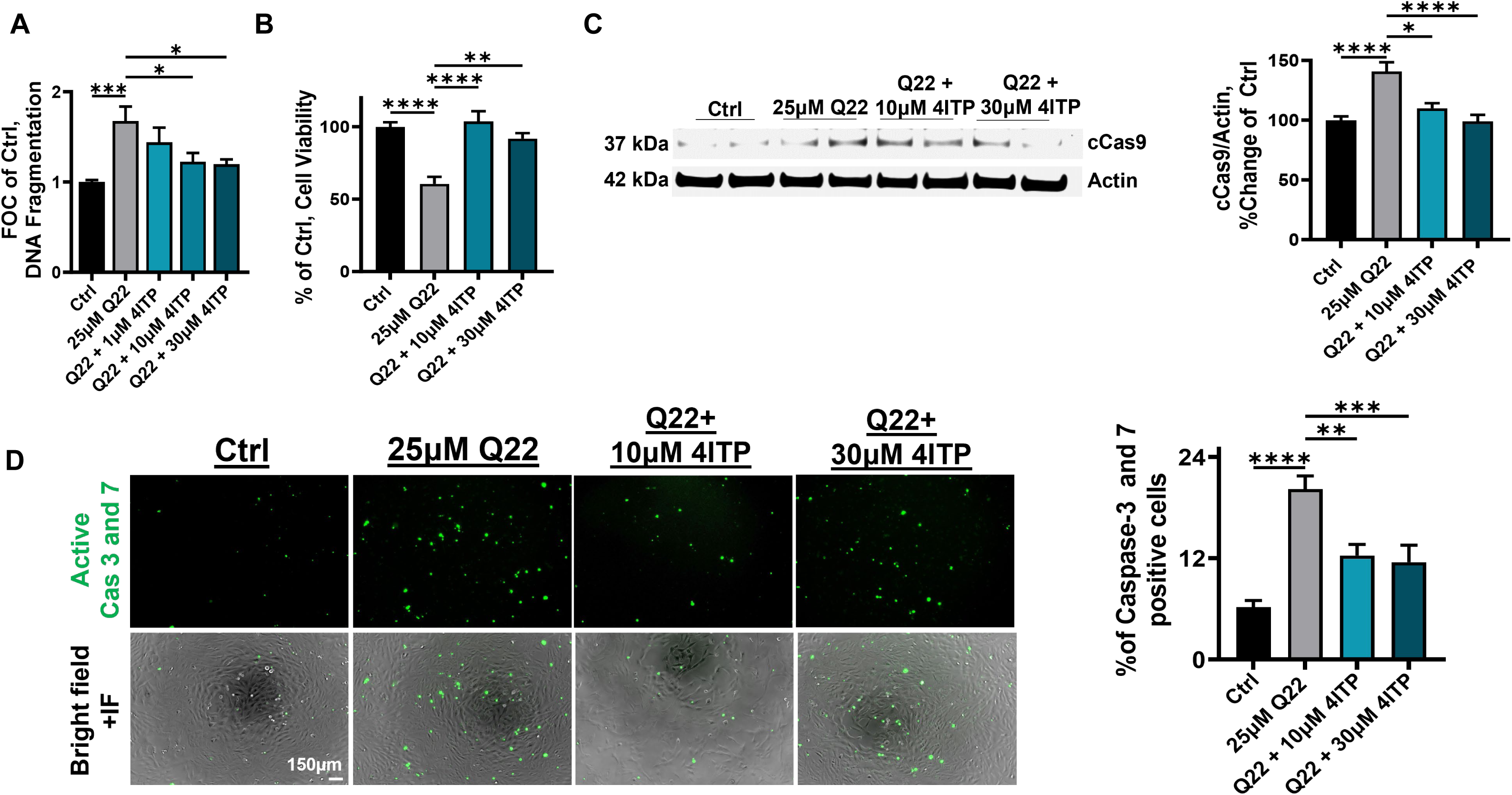
CA-V inhibition protects against Aβ40-Q22-induced apoptosis in hCMEC. 25µM Aβ40-Q22 (Q22) with or without 4ITP (1,10 or 30µM) for 24 h treatment. **A)** Measurement of DNA fragmentation in human CMEC, expressed in fold of change (FOC) of untreated control (Ctrl). **B)** Cell-viability triggered by 24h Q22 treatment, expressed as % of Ctrl. **C)** WB analysis of CMEC lysates to measure cleaved (activated) caspase-9 (cCas9); cCas9 normalized to actin on the right. **D)** Representative IF images of hCMEC, challenged with Q22 alone or in combination with 10 and 30µM 4ITP. Active caspase3/7-positive cells (green) induced by Q22, as plotted on the right. *Original magnification: 10x*. These data represent the combination of at least three experiments, each with 2 replicates, 4 images per replicate, graphed as mean + SEM. Statistical significance was evaluated by One-way ANOVA, followed by Tukey post-hoc test. **** p<0.0001, *** p<0.001, ** p<0.01, * p<0.05

**Figure 2.**
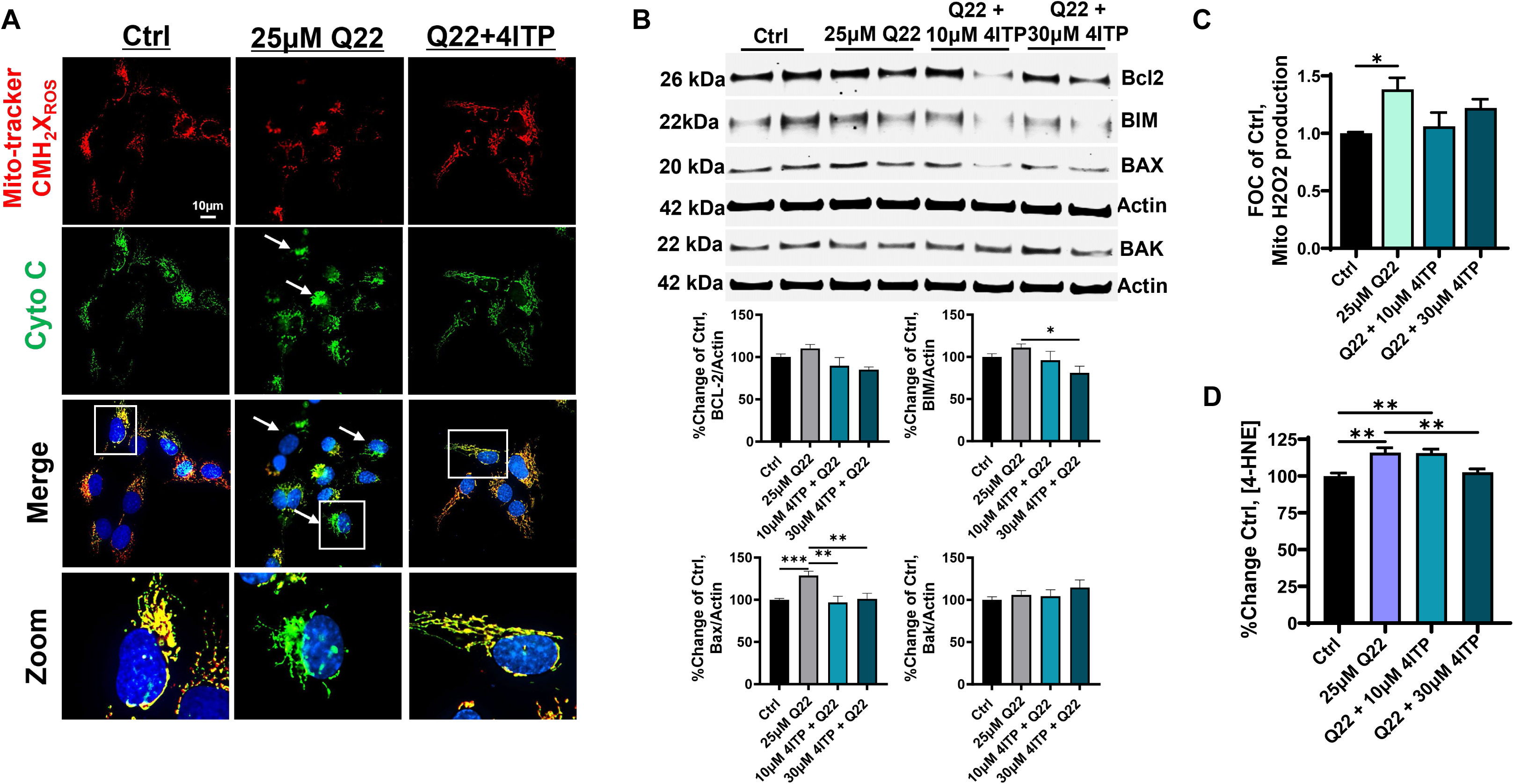
**4ITP protects against Aβ40-Q22 induced mitochondrial apoptotic mechanisms and oxidative stress in hCMEC**. HCMEC treated for 16h with 25µM Aβ40-Q22 (Q22), in the presence or absence of 10 or 30µM 4ITP. **A)** IF images depict the loss of Mito-tracker signal (in red) observed in CMEC challenged with Q22 indicating a loss of mitochondrial membrane potential. The release of cytochrome C (green) is visualized as a diffused, cytosolic signal (indicated with arrows) in Q22-treated cells, compared to untreated cells (ctrl). *Original magnification: 100x.* **B)** Representative WB images of mitochondrial apoptotic mediators BCL-2, BIM, BAX, and BAK, following Q22 treatment, and relative quantification below (loading control, actin). **C)** Mitochondrial H_2_O_2_ production, measured by Amplex Red. **(D)** 4-HNE ELISA, standard curve quantification. Data represent the combination of at least three experiments each with 2 replicates, graphed as mean + SEM. Statistical significance was evaluated by One-way ANOVA followed by Tukey post-hoc test. *** p<0.001, ** p<0.01, * p<0.05

Co-treatment with 10µM 4ITP also prevented AβQ22-induced loss of mitochondrial membrane potential (ΔΨ), measured by the Mito-tracker CMH2xROS dye, which enters only mitochondria with a healthy ΔΨ **(Fig 2A)**. Loss of mitochondrial membrane potential was observed in AβQ22-treated CEC as a decreased Mito-tracker CMH2xROS signal. The release of CytC, measured via immunofluorescence (IF), was evident by a diffused signal in AβQ22-treated cells, indicating CytC release into the cytosol. Both loss of ΔΨ and release of CytC were rescued by 4ITP. Due to these observations, we sought to understand which pro-apoptotic mediators could drive the mitochondrial membrane permeability loss, and if 4ITP could impact their expression. Many proteins may modulate the pore formation on the outer mitochondrial membrane, including pro-apoptotic BAX, BAK and Bim, and anti-apoptotic protein BCL-2 [42, 43]. We observed that the inhibition of CA-V with 4ITP significantly prevented AβQ22-mediated BAX and Bim increase in hCMEC, (**Fig 2B**), while there were no significant changes in Bcl-2 or pro-apoptotic protein BAK expression.

One mechanism that can alter the expression and interactions of mitochondrial pore regulating proteins, as well as affecting ΔΨ, is the accumulation of reactive oxygen species (ROS) [34, 44]. Indeed, after 16h treatment, AβQ22 significantly induced production of H_2_O_2_ by isolated mitochondria, compared to untreated hCMEC. As hypothesized, mitochondrial H_2_O_2_ production was reduced to levels similar to the control by both concentrations of 4ITP (**Fig 2C**). When H_2_O_2_ is produced within the cell, it can react with lipids, forming toxic lipid peroxidation products such as 4-HNE, a known inducer of cell death and marker of oxidative stress, which has also been observed in the brains of AD patients [45]. We measured 4-HNE in CEC treated with AβQ22 and 4ITP. The increase in 4-HNE induced by AβQ22 was significantly prevented by 30µM 4ITP (**Fig 2D**). Overall, these results confirm our hypothesis that inhibition of mitochondrial CA-V prevents Q22-induced mitochondrial ROS production and mitochondria-mediated apoptotic mechanisms.

### CA-V inhibition prevents Aβ driven loss of endothelial barrier integrity

After confirming the positive effects of the CA-V inhibitor on mitochondria-mediated apoptosis and oxidative stress triggered by AβQ22 in hCMEC, we aimed to determine whether CA-V inhibition could also prevent Aβ-mediated loss of endothelial barrier function. To do so, we used a sublethal concentration (10µM) of AβQ22, to observe changes in vascular barrier integrity not due to cell death [20]. The trans-endothelial electrical Resistance (R) of the CEC barrier was measured through the ECIS Z𝜃𝜃 technology, as an indicator of BBB integrity and health, which is decreased in the presence of different Aβ species [20, 21]. Interestingly, co-treatment with 10 and 30µM 4ITP protected against AβQ22-induced loss in R over-time, improving R to levels higher than the control in a dose dependent manner (**Figure 3A**). We also observed that 4ITP alone increased the R compared to untreated control CEC.

**Figure 3.**
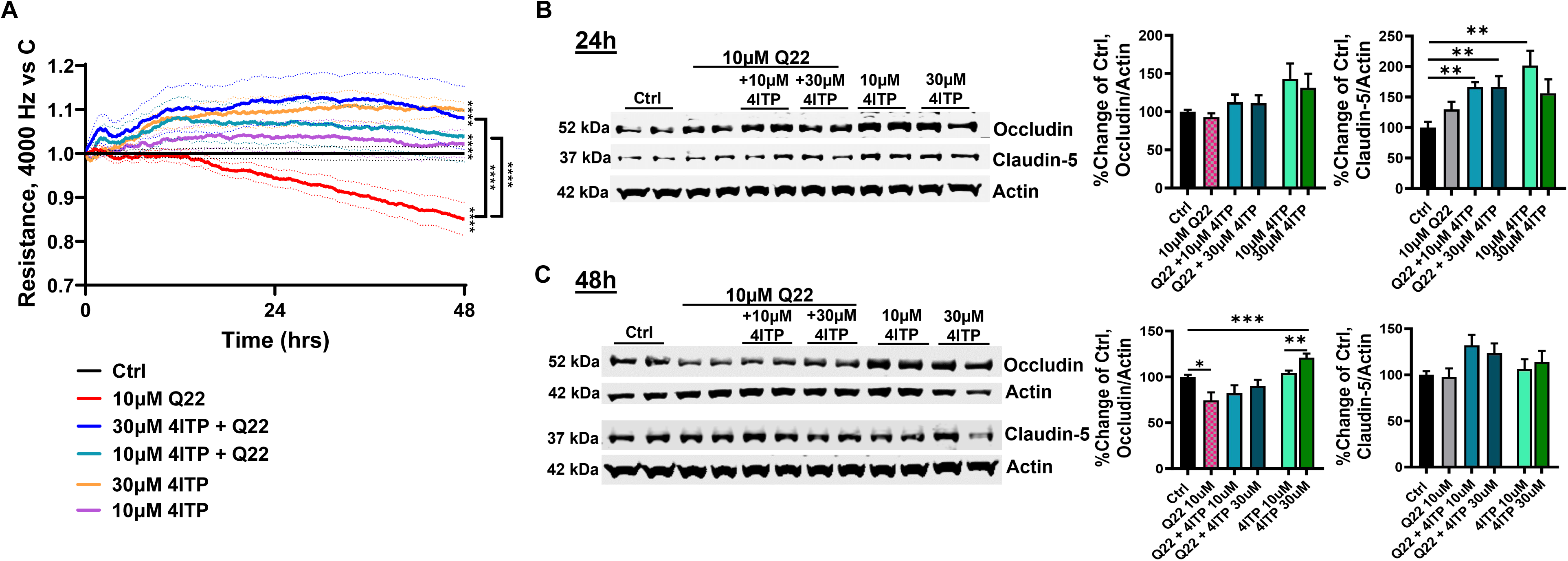
4ITP prevents Aβ40-Q22 induced loss in endothelial barrier integrity. **A)** hCMEC barrier resistance was assessed by ECIS Zθ to evaluate barrier integrity. Human CMEC were treated with 10µM Aβ40-Q22 (Q22), alone or in combination with 10 or 30µM 4ITP. **B-C)** Western blot analysis of occludin and oligomeric claudin-5 (normalized to actin, and plotted as % change of Ctrl), following 24h **(B)** and 48h **(C)** treatment. Data represents 3 individual experiments with 2 replicates per group, graphed as mean ± SEM. Statistical significance was evaluated by One-way ANOVA followed by Tukey post-hoc test. *P< 0.05, **P<0.01, ***P<0.001, ****P<0.0001

To further investigate whether these changes in BBB resistance are associated with modifications in TJP expression, we assessed if 4ITP modulated AβQ22 effects on the expression of TJPs occludin and claudin-5 by WB. At 6h, AβQ22 did not significantly affect the expression of occludin and claudin-5 [46] **(Suppl. Fig 2**), as it did not at 24h (**Figure 3B**). However, 4ITP treatment, alone or in combination with AβQ22, significantly increased claudin-5 expression after 6h **(Suppl. Fig 2)** or 24h **(Figure 3B),** in line with the increased R of CEC monolayers treated with the CA-V inhibitor.

After 48h of treatment with 10µM AβQ22, we observed a significant reduction in the expression of occludin compared to untreated CEC, which was no longer significant in the presence of 4ITP, confirming its protective effects on barrier properties. We also observed a significant increase in occludin expression in cells treated with 30µM 4ITP alone, compared to control CEC **(Figure 3C).** Overall, these results indicate that the CA-V inhibitor 4ITP increases tight junction expression and prevents AβQ22-induced loss of barrier resistance, confirming its therapeutical potential against BBB permeability in cerebrovascular disease.

### CA-V inhibition prevents Aβ-initiated endothelial activation

Another important function of hCMEC is their role as mediators of the crosstalk between the peripheral and CNS immune systems [14]. Due to their bi-directional influence on cerebral and peripheral immune mechanisms, CEC activation state and the production of endothelial inflammatory mediators are critical in AD and other cerebrovascular diseases. To evaluate endothelial activation, we measured the expression of vascular adhesion molecules, known to extravasate peripheral immune cells into the brain, VCAM-1, and ICAM-1, following 6, 24 and 48h of treatment. Interestingly, AβQ22 increased the expression of both VCAM-1 and ICAM-1 at different time-points: at early time points (6h, **Figure 4A)**, ICAM-1 was significantly increased compared to the control, while there were no significant changes in VCAM-1 expression; at later time points (24 and 48h, **Figure 4B and 4C**, respectively), Q22 treatment significantly increased VCAM-1 protein expression, compared to the control hCMEC, while ICAM-1 expression was not significantly changed. Remarkably, co-treatment with 4ITP reduced the AβQ22-mediated increase in ICAM-1 to levels non-significantly different from the control at 6h and significantly restored VCAM-1 levels at both 24 and 48h, in a dose-dependent manner.

**Figure 4.**
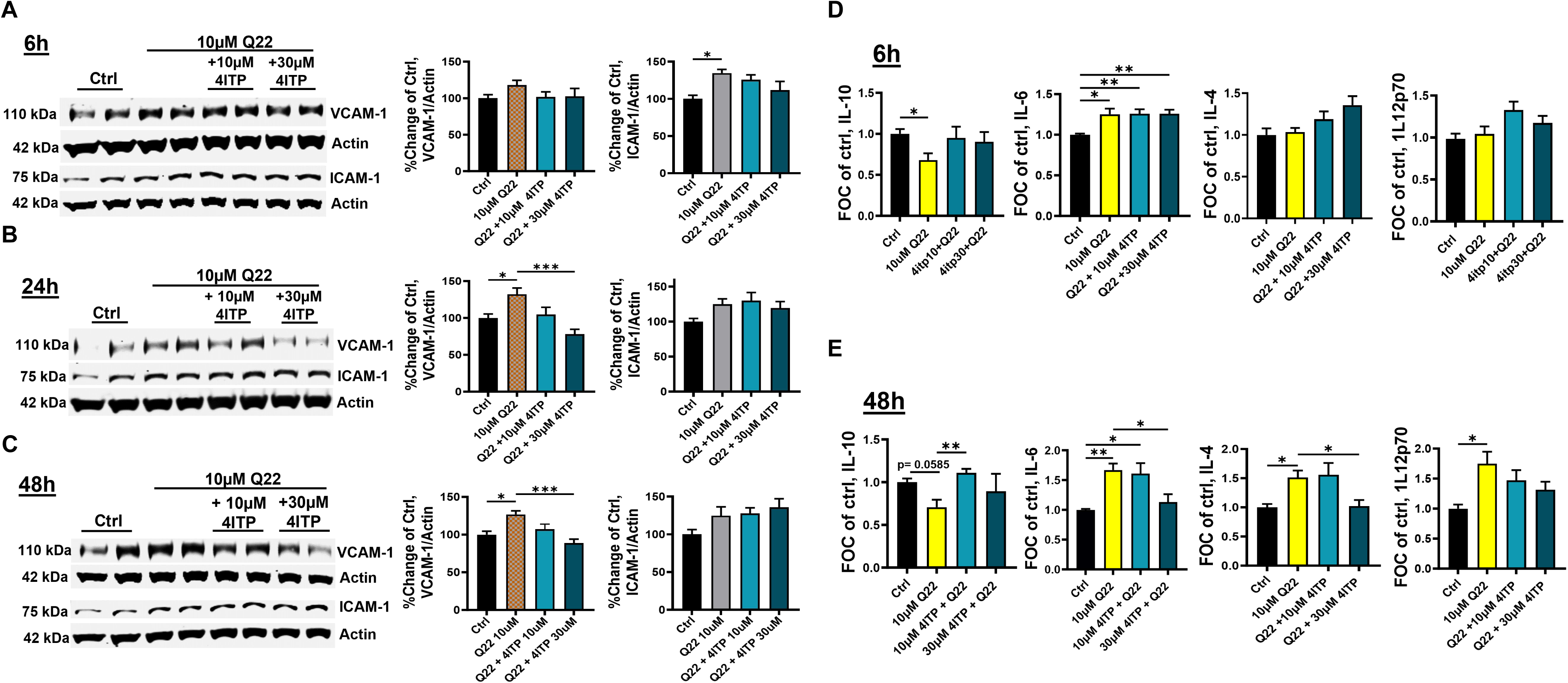
4ITP mitigates Aβ40-Q22 induced brain endothelial activation. Human CMEC were treated with 10 µM Aβ40-Q22 (Q22), alone or in combination with 10 or 30µM of 4ITP. WB analysis of VCAM-1 and ICAM-1 (normalized to actin, and plotted as % change of Ctrl), at **A)** 6h **B)** 24h and **C)** 48h following treatment. **D-E)** Cytokine release was evaluated after 6h and 48h Q22 treatment using a multiplex MSD proinflammatory panel. Data represent 3 individual experiments with 2 replicates per group, graphed as mean ± SEM. Statistical significance was evaluated by One-way ANOVA followed by Tukey’s **(A-C)** or Dunnett’s T3 (**D-E)** post-hoc test. *P< 0.05, **P<0.01, ***P<0.001, ****P<0.0001

We also aimed to correlate the CEC inflammatory mediators released into the conditioned media. Interestingly, after 6h of AβQ22 treatment (**Figure 4D**), we observed a significant decrease in the release of anti-inflammatory interleukin-10 (IL-10), which was reverted by 4ITP. Conversely, we observed a significant increase in the release of pro-inflammatory cytokine IL-6, which was not affected by mitochondrial CA-V inhibition. There were no significant changes in IL-4 or IL-12p70 after 6h of treatment. After 48h **(Figure 4E)**, AβQ22 treatment lowered IL-10 levels (*p= 0.0585*) and increased the release of pro-inflammatory cytokines IL-6 and IL-12p70, and pleiotropic cytokine IL-4, compared to untreated hCMEC. Importantly, 4ITP significantly upregulated anti-inflammatory IL-10 release, compared to Q22 alone (significantly at 10μM), and diminished IL-6, IL-4 (significantly at 30μM), and IL12p70 levels.

To obtain a broader picture of amyloid-induced EC activation we measured mRNA expression of a number of relevant immune cell mediators and cytokines, through a customized Taq-man qPCR array after 24 hours treatment. Confirming the above WB data, we observed a significant Aβ-mediated increase in VCAM-1 mRNA expression which was significantly reduced by 4ITP treatment **(Suppl. Figure 3).** We also observed a significant increase in CXCL2, CSF3/G-CSF, IL-12a/b, IL-4, IL-6, and E-selectin in cells treated with AβQ22. Importantly, CA-V inhibition significantly reduced amyloid-induced increases in CXCL2, CSF3, IL-12b, IL-4, and E-selectin. Oppositely, CA-V inhibition significantly enhanced mRNA expression of anti-inflammatory cytokine IL-10 in the presence of amyloid. Further, we also observed a significant decrease in CXCL1, CSF2/GM-CSF, and P-selectin with 4ITP treatment in the presence of amyloid **(Suppl. Figure 3).** These results suggest that CA-V inhibition ameliorates endothelial inflammatory activation at the transcriptional level, highlighting the therapeutical potential of its inhibition in AD and other cerebrovascular indications.

### CA-VB KO prevents Aβ-induced cerebral endothelial cell apoptosis and loss in barrier resistance

We recently reported that the expression of mitochondrial CA-VB is upregulated in multiple models of amyloidosis, including hCMEC challenged with AβQ22, TgSwDI mice and human AD/CAA brains [25]. We have also demonstrated that silencing CA-VB in hCMEC is protective against Aβ-induced apoptosis. In contrast, expression of CA-VA, the other mitochondrial isoform, or CA-II, the main cytoplasmic CA present in the CNS, did not change. Due to the mounting evidence that CA-VB is likely the mitochondrial isoform mediating Aβ toxicity in AD and to further explore the role of this CA isoform in CEC dysfunction, we used hCMEC CA-VB Crispr/CAS9 KO cells. After confirming almost complete KO of CA-VB **(Figure 5A),** compared to control cells (electroporated with CAS9 alone), we also confirmed that other isoforms of CA, such as CA-VA and CA-II were still expressed in these cells **(Suppl. Figure 4).** Strikingly, after 24h of treatment with 25µM Q22, CA-VB KO hCMEC did not show the significant increase in apoptosis, measured as DNA fragmentation, seen in the WT cells **(Figure 5B)**, suggesting that CA-VB is vital for AβQ22-induced apoptosis in CEC. To further investigate the impact of mitochondrial CA-VB on endothelial apoptosis, we also measured Q22-induced caspase-3/7 activation in CA-VB KO cells **(Figure 5C)**. We found that AβQ22-induced caspase 3/7 activation was completely prevented in CA-VB KO CEC. In addition, IF analysis demonstrated that, after 16h of AβQ22 treatment, AβQ22-mediated loss of mitochondrial membrane potential and cytochrome C release were prevented in CA-VB KO CEC **(Figure 5D)**. Notably, we also observed that CA-VB KO protected from AβQ22-induced mitochondrial H_2_O_2_ production **(Figure 5E)**. Finally, to determine if CA-VB KO protected against barrier integrity loss, we measured endothelial barrier R over-time with ECIS ZΘ. As expected, 10µM AβQ22 triggered loss of barrier R in WT CEC, while in absence of CA-VB, barrier R in the presence of AβQ22 challenge was significantly preserved. A remaining slight loss of R in AβQ22-challenged CA-VB KO CEC monolayers was observed, which may be due to an incomplete KO of CA-VB, or to the involvement of other players, such as CA-VA, which is also inhibited by 4ITP, in Aβ-mediated endothelial barrier dysfunction **(Figure 5F)**.

**Figure 5.**
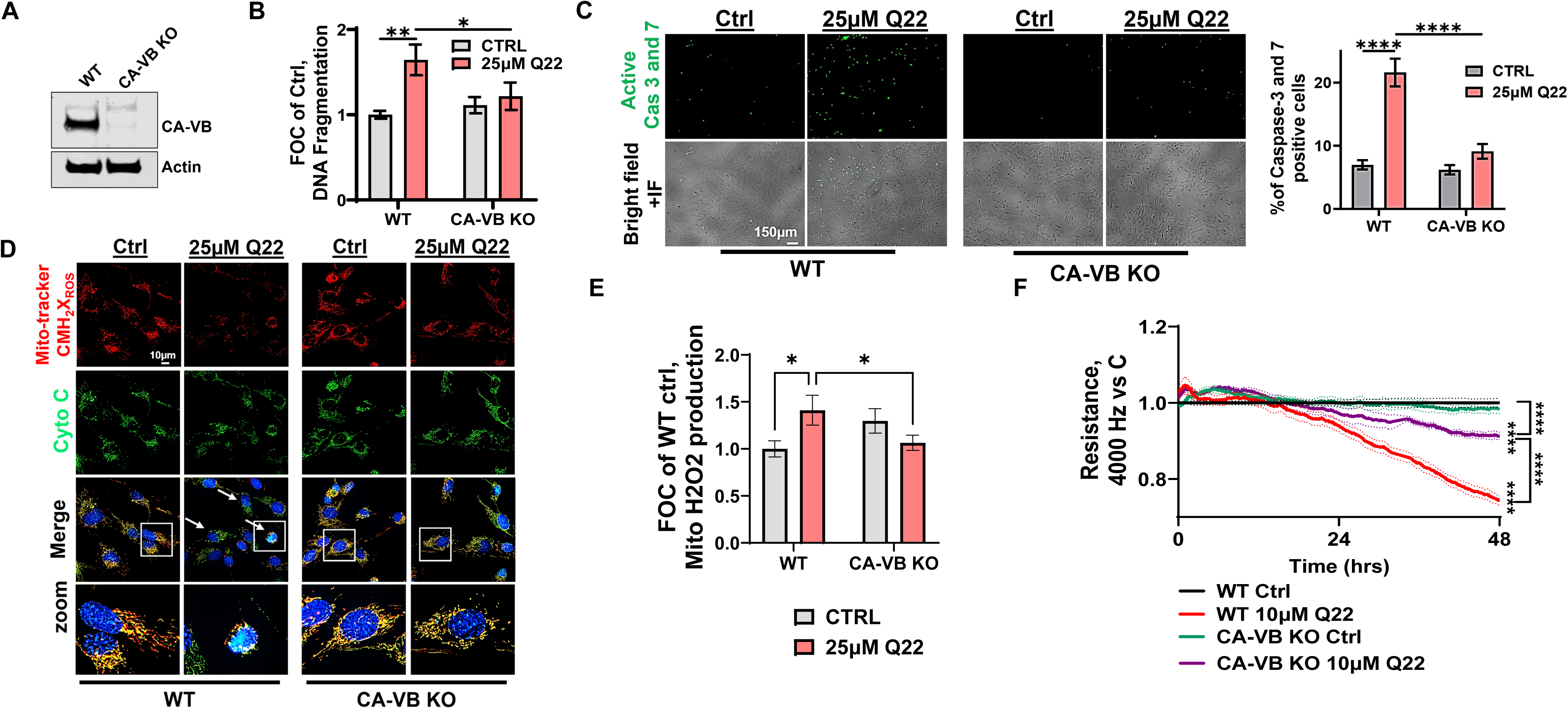
**CA-VB KO in hCMEC protects from Aβ-induced apoptosis and BBB permeability**. **A)** WB confirmed CA-VB absence in KO cells. **B)** DNA fragmentation, plotted as fold of change (FOC) of untreated control (ctrl) cells, measured by cell death ELISA. **C)** Representative IF images of active caspase-3/7 (green) and quantification to the right in CMEC treated with 25µM Q22 for 24 hours. *Original magnification: 10x*. **D)** IF of CA-VB KO vs WT hCMEC to detect mitochondrial membrane potential (Mito-tracker, red) as well as cytochrome C (Cyto C, green), in hCMEC challenged with Q22 for 16 hours. Zoom images of the merge signal at the bottom depict altered mitochondrial network with perinuclear mitochondria in Q22-treated hCMEC, but not in CA-VB KO cells. *Original magnification:100x* **E)** Mitochondrial H_2_O_2_ measured with Amplex Red kit. **F)** Barrier resistance was measured over time with ECIS-Zθ in WT and CA-VB KO hCMEC treated or not with 10µM Q22. Data represents 3 individual experiments with 2 replicates each, graphed as mean ± SEM. Statistical significance was evaluated by Two-way ANOVA followed by Tukey post-hoc test. *P< 0.05, **P<0.01 ****P<0.0001

Due to the complete rescue of amyloid-induced loss in resistance in WT cells treated with 4ITP and incomplete rescue in CA-VB KO cells, we tested whether 4ITP could improve barrier R also in the CA-VB KO cells **(Suppl Figure 5).** We observed that in CA-VB KO hCMEC treated with 4ITP there was still an increase in R, although less than in WT cells. We observed that 4ITP did improve the resistance back to control levels in CA-VB KO cells treated with amyloid. The effect of 4ITP on CA-VB KO cells is not surprising, as this compound inhibits both mitochondrial isoforms, CA-VA and-VB. Overall, these results indicate that CA-VB is necessary for Aβ-induced mitochondrial apoptosis and that its KO significantly ameliorates barrier integrity in CEC monolayers.

### CA-V inhibition in vivo decreases apoptosis and loss of BBB integrity, preventing neuronal loss and cognitive decline

To evaluate the impact of CA-V inhibition *in vivo*, we determined the effects of 4ITP in a mouse model of AD, the 3xTG mouse model, which presents with both Aβ and tau pathology in the hippocampus and cortex. It has been recently demonstrated that these mice also display cerebrovascular deficits and mitochondrial dysfunction [38, 47–50]. Male and female mice (WT and 3xTG) were treated with 4ITP 20 mg/kg/day (delivered through their diet) starting at 6 months, with an endpoint of 16 months. At 6 months of age the 3xTG mouse model is beginning to experience cognitive decline, representing early stages of MCI in humans [41]. The CA-V inhibitor was delivered in their diet. The body weight of treated and non-treated mice was checked weekly, and mice were observed daily to evaluate toxicity throughout the study.

First, we observed that 4ITP treatment in 3xTG mice mitigated overall brain apoptosis, measured as cleaved caspase-3 expression, in both the hippocampus and cortex through immunohistochemistry and WB analysis **(Suppl. Figure 6).** More importantly, we wanted to understand the effects of 4ITP on brain endothelial cell apoptotic mechanisms *in vivo*. Through co-staining of active caspase-3 and CD31, we observed that in the hippocampus of 3xTG mice the % of capase-3 positive vessels was significantly increased compared to WT, while the difference from WT was not significant in the presence of 4ITP treatment **(Figure 6A).** We observed a similar trend in the cortex, although not significant **(Suppl. Figure 7A)**. We also observed a significant reduction of the % area of CD31 in the hippocampus and a similar trend in the cortex of 3xTG mice, suggesting endothelial cell loss. This CEC loss was prevented by 4ITP treatment **(Figure 6B-C).** Additionally, we uncovered a significant decrease in blood vessel diameter in the hippocampus and cortex of 3xTG mice compared to WT. This vasoconstriction was also significantly prevented in the presence of 4ITP treatment **(Figure 6D)**. Overall, these data indicates a preservation in micro-vessels quantity and function in presence of the CA-V inhibitor.

**Figure 6.**
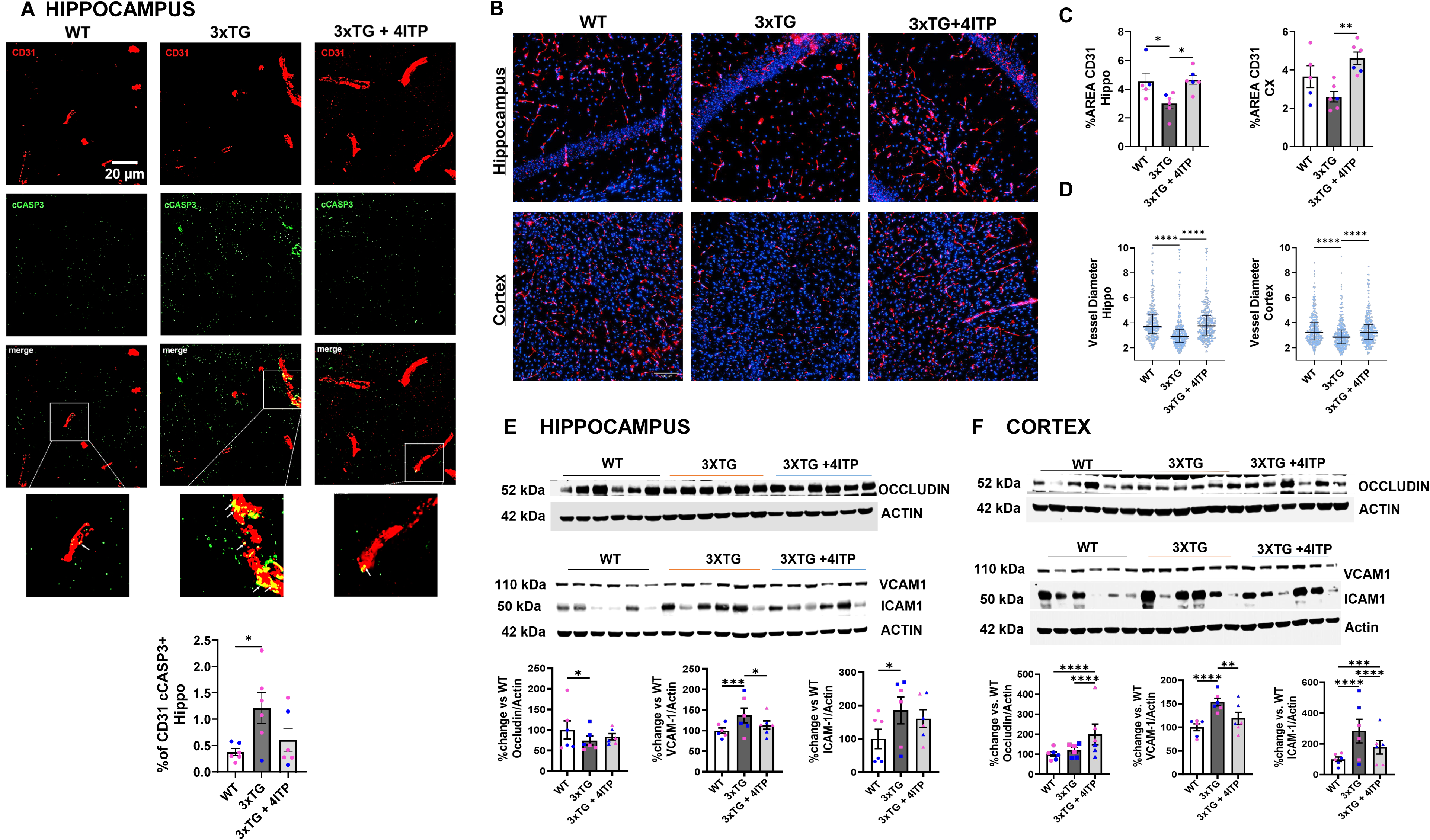
4ITP protects from endothelial cell apoptosis and preserves BBB integrity in 3xTG mice. **(A)** Staining of cCASP3 and CD31 colocalization in the hippocampus of WT, 3xTG, and 3xTG mice treated with 4ITP (as in Methods). CD31 is represented in red and cCASP3 is represented in green. The %Area of colocalized cCAS3 and CD31 was quantified with HALO imaging software. *Original magnification 60x*. Quantification on the right. N=4-7 mice/group, mixed male and female. Data represents 5-10 images per mouse and area. Statistical significance was evaluated by One-way ANOVA followed by Tukey post-hoc test. *P< 0.05, **P<0.01. **(B)** 20x images of WT, 3xTG, and 3xTG + 4ITP cerebral vessels stained with CD31 in red, DAPI in blue. **(C**) Quantification of % area of CD31; **(D)** median blood vessel diameter of WT, untreated 3xTG mice and 3xTG mice treated with 4ITP. **(C-D)** N=5-6 mice/group, mixed male and female. Data represents 5-10 images per mouse and area. Statistical significance was evaluated by One-way ANOVA followed by Tukey post-hoc test. *P< 0.05, **P<0.01 **(E-F)** Western blot analysis of TJP occludin and extravasation proteins VCAM-1, ICAM-1 in the **(E)** hippocampus and **(F)** cortex of WT, 3xTG, and 3xTG +4ITP. **(E, F**) Data represents N=6 mice per group, 2 technical replicates/mouse. Statistical significance was evaluated by Two-way ANOVA followed by Tukey’s post-hoc test (main column effect). *P<0.05, **P<0.01, ***P<0.001, ****P<0.0001. Male and female mice are represented by blue and pink dots, respectively.

We also found that VCAM-1 and ICAM-1 protein expressions were significantly increased in the hippocampus **(Figure 6E)** and cortex **(Figure 6F)** of 3xTG mice, and 4ITP significantly reduced the expression of both these endothelial activation markers in the cortex, and of VCAM in the hippocampus. Finally, in the hippocampus, there was a significant loss in the expression of occludin in 3xTG mice compared to WT, partially mitigated in 4ITP-treated 3xTG mice, which showed no significant difference from the WT animals **(Figure 6E)**. We found no significant changes in the expression of occludin in the cortex between WT and 3xTG mice, although 4ITP induced a significant increase in occludin expression, compared to both WT and untreated 3xTg mice **(Figure 6F).** Our results display some inter-subject variability among some of these markers, particularly ICAM, which appears partially related to sex differences, in line with studies showing high variability in vascular proteins including TJP occludin and extravasation proteins VCAM-1 and ICAM-1 in aging as well as in models of AD [51–54].

To evaluate changes in BBB permeability, we qualitatively assessed albumin leakage in the hippocampus of 3xTG mice. Strikingly, we observed an increase in albumin in the 3xTG mice brains, and particularly, areas of albumin leakage into the parenchyma (indicated with arrows) **(Figure 7A).** These albumin-rich areas were not present in 4ITP treated mice.

**Figure 7.**
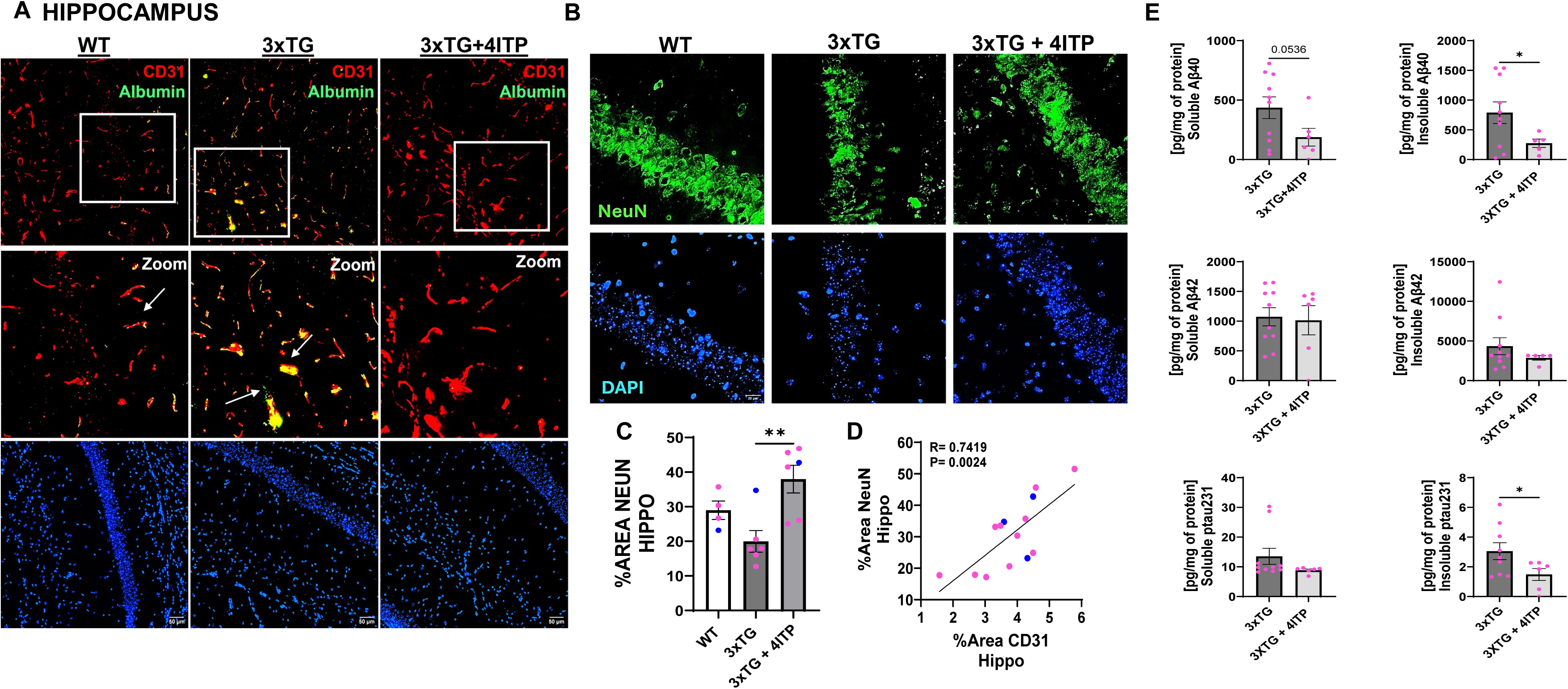
4ITP prevents microhemorrhages, correlating with neuronal and endothelial cell survival, preventing amyloid accumulation. **A)** Representative images of co-staining CD31 and blood protein Albumin (green) in the hippocampus of WT, 3xTG, and 3xTG + 4ITP mice. Microhemorrhages are observed in 3xTG mice (indicated with arrows) and prevented in 3xTG-treated mice. *Original magnification 60x.* **(B)** Representative images of NeuN (green) and DAPI (blue) in the hippocampus of WT, 3xTG, and 3xTG mice treated with 4ITP. *Original magnification 60x.* **(C)** Quantification of %area NeuN in the hippocampus. **(D)** Correlation between %area NeuN and %area CD31 in the hippocampus of WT, 3xTG, and 3xTG + 4ITP mice. 4ITP treated 3xTG had significant improvement in %area of NeuN and the correlation between %area NeuN and %area is significant. **(E)** There is a significant decrease in the insoluble Aβ40 as well as pTau231 in the hippocampus of 3xTG-treated mice. In all graphs, Male and female mice are identified with blue and pink dots, respectively. **(C-D**) N=4-6/group (indicated with dots). Statistical significance was evaluated by One-way ANOVA followed by Tukey’s multiple comparison test. **E)** N=10 female 3xTG N=6 female 3xTG treated. Statistical significance was evaluated with T-test Welch’s correction between 3xTG and 3xTG-treated mice. *P< 0.05, **P<0.01 ****P<0.0001.

To have a better understanding of the effects on neuroinflammation *in vivo* and continue to validate our *in vitro* findings, we measured cytokines and chemokines in the cortex of WT, 3xTG and 3xTG 4ITP-treated mice. Similarly to the *in vitro* data, we observed a trending decrease in anti-inflammatory cytokine IL-10, which was significantly prevented in the brains of 4ITP treated mice. We did not observe changes in IL-6 or IL-12p70 in the cortex of these mice. Differently, we saw a decrease in pleiotropic cytokine IL-4 in brains of 3xTG mice which was significantly prevented in 4ITP treated mice (**Suppl. Figure 7B**). We also observed trends in increases in KC-GRO/CXCL1 as well as MIP-2/CXCL2 in 3xTG cortex, which were mitigated in 4ITP treated 3xTG mice.

As neuroimmune cell activation may be associated to endothelial cell death as well as endothelial cell activation, we measured the extent of astrogliosis in 4ITP treated 3xTG mice **(Suppl. Figure 8A)**; in the hippocampus and cortex, the % area of GFAP was significantly reduced in mice treated with 4ITP compared to untreated 3xTG mice. We also measured the concentration of 4-Hydroxynonenal (4-HNE), a highly reactive aldehyde produced during lipid peroxidation, which is considered a marker of oxidative stress, in the cortex of WT, 3xTG non-treated and treated mice, but we did not detect any significant changes **(Suppl. Figure 8B).**

To understand if the prevention of endothelial cell loss and preservation of BBB integrity are associated to the deterrence of neuronal loss and progression of neurodegenerative disease, we measured the % area of NeuN, a neuronal cell marker. We observed a decrease in NeuN reduction in the hippocampus of 3xTG mice, and the % area of NeuN was significantly increased in the hippocampus of 4ITP treated mice compared to 3xTG mice **(Figure 7B-C).** There were no significant changes in NeuN % area in the cortex of these mice **(Suppl. Figure 9A-B).** To understand the relationship between the endothelial cell and neuronal cell loss in these mice, we measured the correlation between the % area of NeuN+ to the % area of CD31+ cells in the hippocampus **(Figure 7D)** and cortex **(Suppl. Figure 9C)** of WT, untreated 3xTG, and treated 3xTG mice. Intriguingly, we revealed a positive correlation both in the hippocampus *(R=0.74, p=0.0024)* and in the cortex (R=0.5308, p=0.0508).

Further, we measured the levels of amyloid and phosphorylated tau in the hippocampus and prefrontal cortex of 3xTG female mice (as male 3xTG mice are known to have minimal accumulation of amyloid or P-tau at this age [41, 55]). In the hippocampus, soluble and insoluble vasculotropic Aβ40 was significantly reduced in 4ITP-treated mice compared to untreated, suggesting a reduction in CAA, as we have previously shown for pan-CAIs [25]. Insoluble ptau-231 was also significantly reduced in the hippocampus of 4ITP-treated 3xTG mice **(Figure 7E).** Although there was much less accumulation, the trends were similar in the prefrontal cortex, where we observed a significant reduction in soluble and insoluble Aβ42, and a trending reduction in soluble Aβ40, while insoluble Aβ40 was not detected. Finally, there were no significant changes in ptau-231 in the prefrontal cortex of treated or untreated 3xTG mice **(Suppl. Figure 9D).**

We then asked whether these beneficial changes in the brains of 4ITP-treated 3xTG mice were correlated with an improvement in cognition. To achieve this goal, we evaluated the mice’s cognitive ability over a 6-day period using the Barnes Maze, a paradigm employed to measure the ability to consolidate and retrieve spatial memories. As expected, 3xTG mice exhibited an increased latency to find the escape hole throughout the learning trials. We first tested two different treatment times (6-16 months and 12-16 months) in the Barnes Maze, and the long-term 4ITP treatment (6-16M), akin to treating MCI subjects in human trials, was much more successful at improving spatial memory compared to the shorter treatment **(Suppl. Figure 10)**. Therefore, we focused on the longer (6-16M) treatment throughout this study.

A significant difference between treated (6-16M) and untreated mice was observed on day 1 and 2 (training) of the maze **(Figure 8A)**. Moreover, 4ITP-treated 3xTG mice did not show significant differences from WT mice on any day. On probe day (day 6), untreated 3xTG mice demonstrated a significant increase in the time to find the escape hole (latency) and number of mistakes, which was significantly reduced to WT levels in 4ITP-treated 3xTG animals (**Figure 8B).** Additionally, we did not observe any spatial memory differences between WT and WT + 4ITP (6-16M) mice, suggesting lack of undesired cognitive effects of 4ITP in heathy subjects **(Suppl. Figure 11A-B).**

**Figure 8.**
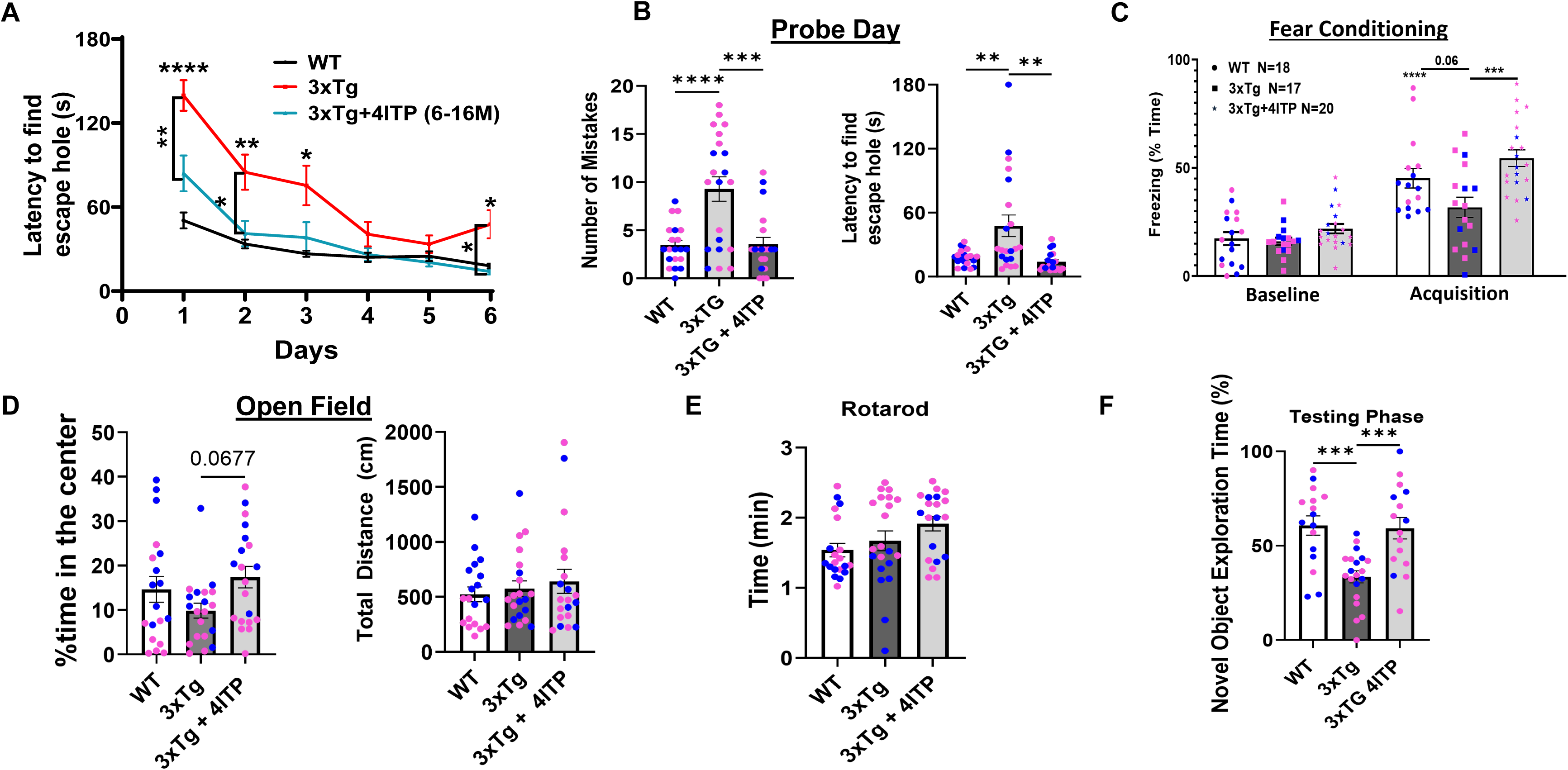
4ITP prevents cognitive decline in 3xTG mice. (WT, N=20 3XTG, N=21 3XTG+4ITP N=20, males, and females 16 months) Spatial memory was assessed using Barnes Maze over a 6-day period **(A)** WT, 3xTG treated and non-treated mice learning trajectory (latency, s) to find the escape hole over-time. **B)** On probe day, the number of mistakes *(F=10.86(2,58) p<0.001)* and latency to find the escape hole *(F=7.173(2,57) p=0.0017)* in each group was recorded. **C)** Cued fear-conditioning test is improved in 3xTG mice treated with 4ITP. **D)** Open field total distance travelled (motor activity) displayed no significant differences. The % of time in the center showed a trend to increased time in the center (decreased anxiety) for 4ITP-treated 3xTG mice compared to untreated 3xTg. **E)** Rotarod test displayed no locomotor differences between groups. **F)** Novel object recognition test is impaired in 3xTG mice and then improved in mice treated with 4ITP. Male and female mice are identified with blue and pink dots, respectively. **A)** Statistical significance was evaluated by Two-way ANOVA followed by Tukey’s multiple comparison test. **B-F)** Statistical significance was evaluated by One-way ANOVA followed by Tukey post-hoc test. *P< 0.05, ** P<0.01 ****P<0.0001.

In a fear-conditioning paradigm, assessing cue-induced fear associative memory, we observed a memory impairment (reduced freezing time) in 3xTG mice compared to WT, which was significantly reverted in 4ITP-treated mice **(Figure 8C)**. We did not see an impairment in WT treated mice, again indicating a lack of undesired side effects **(Suppl. Figure 11C)**. Through open field test and rotarod, we detected no significant locomotor changes between wildtype and 3xTG mice **(Figure 8D-F)** or WT and WT + 4ITP **(Suppl. Figure 11D and 11E).** Additionally, in the open field test, 4ITP-treated 3xTG mice showed a trend to increased time in the center compared to untreated 3xTG animals, suggesting a reduction in anxiety behavior upon treatment. Finally, we observed a preservation in object memory of 3xTG treated mice. With the novel object recognition test, we observed a significant reduction in the % time exploring the novel object during the testing phase in 3xTG mice compared to WT, indicating a loss in working memory. The time exploring the novel object was significantly increased in 4ITP treated 3xTG mice, indicating a preservation of working memory **(Figure 8F).**

When separating the mice by sex we did observe some sex differences, although not significant **(Suppl. Figure 12)**. Notably, 3xTG male mice were larger and moved slower than female mice in the Barnes maze, as indicated by their similar trends but different degrees of significance in the latency to find the escape hole **(Suppl Figure 12A-B and 12D-E)**, as previously reported [53, 55]. However, the latency to find the escape hole on probe day was improved by 4ITP in both males and females. Both males and female 3xTG mice made a significantly higher number of mistakes compared to wild-type mice on probe day, but 4ITP treatment resulted in a more significant improvement in the number of mistakes in female mice **(Suppl Figure 12B and E)**. Additionally, we observed a rescue in cue-associated fear memory in both females and males **(Suppl. Figure 12C and G).** In the novel object recognition test, we observed an increase in %time exploring the novel object in both female and male 4ITP-treated mice; however, this increase was significant in male treated mice **(Suppl. Figure 12D and H).** We did not observe any significant differences in the open-field test when analyzing the % of time spent in the center, or the total distance travelled **(Suppl. Figure 12I-J)**. Interestingly, there was a significant improvement in motor function (rotarod) in male 4ITP-treated 3xTG mice compared to untreated 3xTG mice **(Suppl. Figure 12K).**

Overall, these data confirms that CA-V inhibition, by preventing apoptosis, cerebrovascular dysfunction, and neuroinflammation in the brain of 3xTG AD mice, significantly rescues cognitive impairment.

## Discussion

This study demonstrates that mitochondrial CA-V is a promising target for the treatment of AD, CAA and related cerebrovascular pathologies. We showed that the selective CA-V inhibitor 4ITP, more lipophilic and effective on CA-V than ATZ and MTZ, prevents endothelial Aβ-induced mitochondria-mediated apoptosis, loss of mitochondrial membrane potential, accumulation of oxidative stress, loss of hCMEC barrier resistance, and endothelial inflammatory activation. Additionally, through CA-VB CRISPR-KO cells, we confirmed mechanistically that the mitochondrial isoform CA-VB is necessary to promote AβQ22 induced endothelial mitochondria-mediated apoptosis, oxidative stress and loss of EC barrier integrity. Strikingly, the same CA-V inhibitor was protective in the 3xTG mouse model through the prevention of apoptosis, cerebrovascular dysfunction and inflammatory activation, resulting in a reduction in AD pathology and a potent improvement in memory.

Cerebrovascular and mitochondrial dysfunction are early and causative events observed in many diseases associated with aging [7, 9, 10, 14, 20, 26, 56–58]. Both cerebrovascular and mitochondrial dysfunction are observed to occur even prior to Aβ and tau accumulation, and worsen throughout the progression of AD and CAA, causing a vicious loop of impaired clearance, toxic protein accumulation, neuroinflammation, exacerbated cerebrovascular and mitochondrial/metabolic deficits. Due to the high impact of cerebrovascular and mitochondrial deficits on the development and progression of AD and CAA, it is evident that these are both ideal targets for the development of novel therapeutic strategies against AD and CAA.

Current FDA-approved therapies for AD do not appear to improve cerebrovascular function. On the contrary, toxic effects of the new anti-Aβ antibodies, increasing Aβ deposits on the BBB, are likely to be responsible for the development of ARIA-E (edema) or ARIA-H (hemorrhages) in a substantial percentage of patients treated with these new drugs [5, 59], emphasizing the importance and need for treatments targeting vascular dysfunction in AD and CAA. Moreover, mitochondrial and metabolic deficits are also frequently discussed among the main pathways to target for the development of new drugs against AD and dementias. However, mitochondria-targeted therapies have not yet reached FDA approval for these diseases [14, 57, 58]. Importantly, a multitude of evidence demonstrates that lifestyle factors such as diet and exercise, as well as cardiovascular risk factors, can influence both BBB and mitochondrial function, resulting in the modulation of risk and progression of symptoms associated with the disease [1, 7, 60, 61]. Cardiovascular risk factors, in particular, may increase cerebrovascular complications in AD and CAA patients, through their toxic effects on CEC [7, 21].

CAs catalyze the hydrolysis of carbon dioxide to produce bicarbonate and a proton. This is an essential reaction to maintain physiological cell health, and particularly within the mitochondria, due to the high production of carbon dioxide as a by-product of metabolism. The use of CA inhibitors has been first explored in cardiovascular disease [62, 63]. Pan-CA inhibitors such as ATZ and MTZ were developed as legacy diuretics, being optimized to modulate the function on the kidney and water reabsorption. They are relatively polar (log P ATZ =-0.26, log P MTZ = 0.13) as they were optimized to work primarily at the level of the kidneys. They have then become FDA-approved for many diseases such as glaucoma, epilepsy, and high-altitude sickness [2, 34]. More recently, they have also been demonstrated to be effective in models of obesity, cerebral edema, stroke, and diabetes-induced cerebrovascular pathology [27, 30, 64]. Many of the studies, dissecting the involvement of CA in cardiovascular pathology or stroke, have used pan-CA inhibitors, most commonly ATZ and MTZ [2, 65]. As previously mentioned, ATZ and MTZ have also been proved effective against Aβ-mediated toxicity in many NVU cells [23, 24, 26], and in a CAA mouse model, the TgSwDI [25]. Additionally, topiramate, another CA inhibitor, has been demonstrated to be protective in models of diabetes-induced cerebrovascular pathology [64, 66]. Although many pan-CA inhibitors have protective effects, they are not ideal for therapeutic use, as their lack of selectivity requires significantly higher doses to achieve efficacy, which cause undesired side effects when used systemically in humans [67].

Our previous studies have demonstrated that ATZ and MTZ prevent Aβ toxicity, specifically through mitochondria-mediated mechanisms [19, 23, 24, 26]. Here, we tested for the first time a mitochondria-selective CA-V inhibitor, and we showed that it is effective at similar concentrations to ATZ (the most effective of the pan-CAIs we have tested *in vitro* and *in vivo*), against mitochondria-mediated apoptosis in hCMEC [24]. Hence, we anticipate that selective CA-V inhibitors will be at least as effective as pan-CA inhibitors against AD/CAA, while avoiding side effects that have been reported in humans for pan-CAIs. One of the possible side-effects to be considered when using high concentrations of pan-CA inhibitors, such as ATZ, is short-term memory impairment. Indeed, a high dose of ATZ has been observed to inhibit long-term potentiation, acutely, in mice [68, 69]. Additionally, humans have reported memory reduction when taking ATZ at high concentrations [68]. This effect may be due to the activity of non-selective inhibitors on a specific CA isoform, CA-VII, abundantly expressed in pyramidal neurons of the hippocampus [70]. However, the dose of ATZ used in our animal models (20mg/kg/day), which is the same dose of 4ITP in the present study, corresponds to a human dose of 2.6 mg/kg/day (182 mg/day for a 70 kg person), and is lower than the FDA-approved dose for CAIs (ATZ: up to 1g/day), which is the dose known to induce short-term memory issues. Nevertheless, to confirm the safety and lack of this side effect for 4ITP, we have shown here that, while 4ITP treatment ameliorates memory in the 3xTG AD mouse model, the chronic use of 4ITP on WT animals at the proposed concentration does not impair spatial memory. Additionally, when comparing two courses of treatment, starting at 6 or 12 months, we observed that the long treatment was much more effective than the later treatment. This results suggests CA-V inhibition is likely targeting early mechanisms of the disease such as cerebrovascular and mitochondrial dysfunction as well as neuroinflammation.

Due to the diverse expression pattern and functions of CAs, it is evident that not all 15 CA isoforms are involved in the development of AD pathology. CA-V, expressed in the mitochondrial matrix, has been demonstrated to influence intermediates of the citric acid cycle through the supply of bicarbonate for enzymes like pyruvate carboxylase and propionyl carboxylase, thereby affecting biosynthetic pathways, such as gluconeogenesis and ureagenesis. However, many of these studies have been done in the liver, where CA-VA is much more abundant. Interestingly, the involvement of CA-VB in metabolism is only evident in the absence of CA-VA [71]. This is observed in cells where CA-VA is not expressed, as well as in Car5a KO mice. Less is known about the function of CA-VB; however, it is much more abundant throughout the body and in the brain [2]. Interestingly, Car5B global KO mice, differently from Car5A global KO mice, are viable and show normal growth and normal ammonia levels, suggesting that chronic inhibition of CA-VB or Car5B KO does not have toxic effects [71].

We have shown here that knocking out CA-VB completely prevents mitochondria-mediated apoptosis induced by Aβ in hCMEC and mitigates the loss of barrier resistance. Thus, selectively targeting the mitochondrial isoform CA-VB, while avoiding undesired effects on other CA isoforms not involved in the progression of AD and CAA, may be of high interest for therapeutic development in AD related disorders, as well as, potentially, against cerebrovascular complications induced by current treatments with FDA-approved anti-Aβ antibodies (such as ARIA).

Interestingly, while 4ITP, which inhibits both CA-VA and-VB, completely rescues barrier resistance in presence of Aβ, CA-VB KO only partially mitigates Aβ-induced BBB permeability. When treating CA-VB KO cells with 4ITP however, we do see a complete rescue in resistance. These constitutes important knowledge, as 100% selectivity for CA-VB is not easily achievable, and these results suggest that CA-VA (or additional CA isoforms 4ITP may inhibit) could also be involved in the induction of BBB permeability in CEC. It is clear however, that 4ITP, as well as CA-VB KO, do significantly improve barrier resistance and should be further explored as a potential therapies against cerebrovascular pathologies. Future studies comparing CA-V inhibitors and their selectivity profiles would help us to better understand the other players that may be involved. We are confident that 4ITP is selective for CA-V compared to other highly expressed CA isoforms [36], however, other, even more selective CA-VB inhibitors will be compared in future studies to further investigate the pharmacodynamics of CA-V inhibition.

Besides CA-V’s role in metabolism, we have demonstrated here that CA-VB activity directly influences mitochondria-mediated apoptosis and mitochondrial membrane potential, as well as oxidative stress. We anticipate that, while CA-VA may function mainly to regulate metabolism, CA-VB regulates mitochondrial pH and redox homeostasis. Mechanistically, the inhibition of CA activity in the mitochondrial matrix is expected to reduce the over-production of hydrogen ions, especially in times of stress (such as amyloidosis), maintaining a lower proton level within the mitochondrial matrix compared to the intermembrane space. A lower H^+^ concentration in the matrix will aid in preserving a healthy mitochondrial membrane potential, in addition to an optimal proton motive force, especially in the presence of ROS. This is essential for proper mitochondrial function, cell signaling and, ultimately, cell survival. Indeed, we have demonstrated that 4ITP prevents the loss of mitochondrial membrane potential, mitochondrial H_2_O_2_ production, as well as CytC release, confirming the mitochondria-specific mechanism of action of the drug, as also supported by the same results in CA-VB KO cells.

A plethora of evidence indicates that preservation of mitochondrial function and mitigation of ROS prevents downstream neuroinflammatory mechanisms. ROS production and release have been observed to induce pro-inflammatory cytokine production, resulting in the activation of glial and vascular cell types. In hCMEC and mouse cortex we observed a preservation in anti-inflammatory cytokine IL-10 with 4ITP treatment. The expression of IL-10 has been observed to negatively correlate with mitochondrial function as well as oxidative stress [72]. This is an indicator that the prevention of ROS through 4ITP could be preventing the downregulation of IL-10, allowing for improved cellular function and cell survival, as well as diminishing chronic inflammatory mechanisms observed to cause damage in AD.

When CEC are activated, they increase their expression of adhesion proteins VCAM-1, E-selectin, P-selectin and ICAM-1 as well as releasing chemokines and cytokines CXCL1/2 and CSF2/3 known to promote immune cell extravasation and EC activation [51]. While recruitment of immune cells is important for the clearance of toxic substances from the brain, chronic inflammation and CEC activation is damaging to the BBB and thus to the NVU and the brain. The expression of VCAM-1 was decreased by 4ITP in the presence of AD pathology. CXCL2/MIP-2, a known recruiter of neutrophils, was also mitigated by 4ITP in vitro and in vivo. Neutrophils have been observed to be very damaging to brain tissue, particularly through the injury of capillaries [73, 74]. BBB dysfunction and EC activation lead to many downstream negative consequences, exacerbating a viscous cycle of cerebrovascular dysfunction and neuroinflammation, including glial cell activation [2, 14, 25], and eventually neuronal loss. Intriguingly, the observed prevention of CEC activation and preservation of endothelial cell function as well as mitigation of astrogliosis correlate with a preserved number of hippocampal neurons, and dramatically improved cognition in 4ITP treated 3xTG mice.

The accumulation of Aβ around the brain vessels, found in AD, sporadic CAA, CAA related inflammation (CAAri), as well as in ARIA complications due to high amounts of Aβ unable to be cleared by the cerebral vasculature, is known to exacerbate BBB dysfunction and induce permeability, enhancing the likelihood of ischemic and hemorrhagic strokes and white matter abnormalities [6, 75]. Therefore, we anticipate that mitochondrial CA inhibitors could not only be a potential therapy for AD and CAA when used alone, but that the combination of CA inhibitors and anti-Aβ therapies could be a very beneficial co-treatment strategy. Further studies are needed to confirm this hypothesis. Interestingly, we observed reduction of amyloid and phosphorylated tau accumulation in the hippocampus of 3xTG-treated female mice, likely due to preservation of cerebrovascular function and reduction of neuroinflammation, which facilitate improved clearance. Importantly, we have demonstrated that both female and male mice are experiencing vascular abnormalities, endothelial cell activation, and gliosis. Notably, 4ITP also rescues cognition in both females and males **(Figure 8 Suppl. Figure 12**). These results suggest that CA-V inhibition is protecting against the toxic effects of amyloid and, more strikingly, against the underlying mechanisms amyloid exacerbates, such as endothelial cell stress/apoptosis, endothelial and neuronal cell loss, and neuroinflammation, getting to the root causes of this multi-factorial disease through multiple of the mechanisms that are known to cause it or mediate it.

One limitation of this study is that it is currently impossible to have complete pharmacological selectivity for one CA-V isoform. 4ITP has been previously shown to have a Ki value of 8.6 nm and 8.3 nm for CA-VA and-VB respectively, indicating equal selectivity for the mitochondrial isoforms. 4ITP is about 8x less effective on CA-I and-II (Ki = 75 and 54 nm respectively), and even less on plasma membrane isoforms CA-IX and CA-XII (136 and 212 nm) [35, 36]. Although this study, by comparing the effects of 4ITP with CA-VB KO cells, provides strong evidence that 4ITP effects have CA-V specificity, the activity of 4ITP on other CA isoforms relevant to the brain/NVU activity remains to be determined. Of interest, CA-IV is highly expressed on the plasma membrane of CEC and should be considered when studying CA function in CEC. Furthermore, knowledge on the activity of 4ITP on CA-VII would be informative, as this isoform is involved in memory and highly expressed in neurons. The lack of an effect of 4ITP on WT animals’ behavior, however, suggests low affinity for this isoform. Studies providing more evidence about the selectivity of 4ITP would help us understand the mechanism of action and to design other selective CA-V inhibitors in future studies.

Another limitation is that we did not measure BBB function or CBF in real time *in vivo*. However, we did uncover a decrease in blood vessel diameter in the brains of 3xTG mice compared to WT. Importantly, this was rescued with 4ITP suggesting it did in fact improve vessel function and integrity. We also showed a prevention of albumin leakage into the brain, suggesting reduced BBB permeability *in vivo*. Future studies under development in our laboratory intend to assess the effects of CA-V inhibitors and cerebral endothelial-specific CA-VB KO, on CBF, BBB permeability and neurovascular coupling in AD mouse models. These studies will also determine the effects of 4ITP on oxidative stress, microglia phenotype and neurodegeneration markers in vivo, as we have previously shown for MTZ and ATZ [25]. We did not focus on the effects of 4ITP on microglia in this manuscript, as this is an ongoing study in the lab and will be published with detail in the future. The strong protective effects of 4ITP on CEC function and on cognition shown here suggest potential benefits in all those mechanisms.

Co-culture studies would help us further understand the direct connection between amyloid induced endothelial activation and neuronal cell survival. The lack of co-culture experiments is a limitation of this study, and future studies could employ co-culture systems to determine if the inhibition of CA-V in endothelial cells treated with amyloid can prevent direct neuronal toxicity.

Additionally, only post-mortem measures of oxidative stress with 4-HNE were performed in this study. Future studies will be needed to assess mitochondrial function, metabolism, and oxidative stress outcomes in fresh brain tissue by seahorse assays of metabolic/mitochondrial function, single cell analysis or other methods.

Overall, this study is pioneer in demonstrating the protective effects of selective CA-V inhibition in models of amyloidosis and AD. We have shown that the CA-V inhibitor 4ITP, as well as CA-VB CRISPR KO, protect CEC from Aβ-induced mitochondria-mediated apoptosis and BBB dysfunction, and that 4ITP is effective to reduce brain apoptosis, endothelial cell survival, cerebrovascular and neuroimmune function, and cognitive impairment in 3xTG mice. Taken together, it is evident that the development and testing of CA-V inhibitors and understanding the role of mitochondrial CA-VB are critical areas of research for therapeutic targeting of cerebrovascular pathology and mitochondrial dysfunction in AD, CAA, ADRD and the vascular contributions to cognitive impairment and dementias.

## Acknowledgments

Main figures were created with BioRender.com (Figures 1-8). Created in BioRender. Fossati, S. (2025) https://BioRender.com/zg3f90e

## Funding

This work was supported by: NIH R01NS104127 (SF) NIH R01AG062572 (SF)

Edward N. and Della L. Thome Memorial Foundation Awards Program in Alzheimer’s Disease Drug Discovery Research, (SF)

The Alzheimer’s Association (AARG) (SF)

Pennsylvania Department of Health Collaborative Research on Alzheimer’s Disease (PA Cure) Grant (SF)

Karen Toffler Charitable Trust (SF and NL)

## Author contributions

Conceptualization and study design: NLL, MAI, SF Methodology: NLL, SF, MAI

Experiment for manuscript: NLL, LMP, TR, EC, RVT, RMP, TH, MHA Supervision: SF, MAI

Writing—original draft: NLL, SF, EC

Writing—review & editing: NLL, EC, RVT, RMP, TH, LMP, TR, MAI, SF

