## Supplemental Figures for "Mitochondrial carbonic anhydrase-VB inhibition rescues brain endothelial stress and memory in Alzheimer’s disease models"

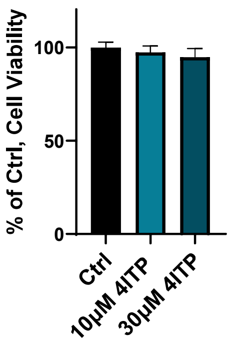


**Figure S1. 4ITP is not toxic at concentrations of 10 or 30µM in hCMEC.** There is no significant change in cell-viability, plotted as % of Ctrl, after 24-hour 4ITP treatment.

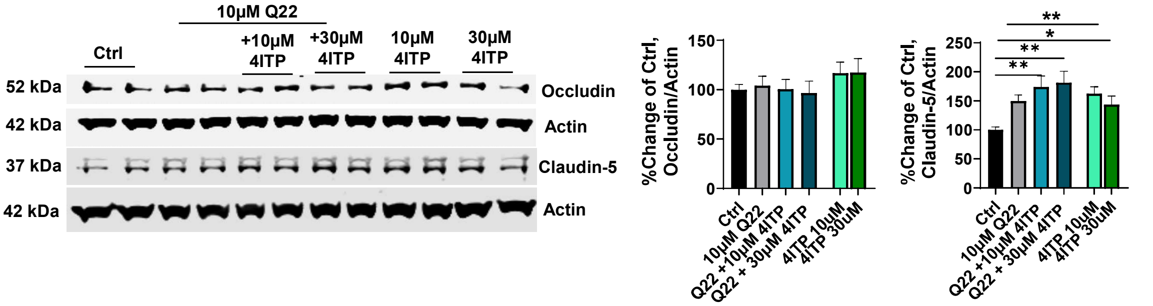


**Figure S2. 4ITP increases expression of claudin-5 in hCMEC after 6 hours of treatment.** Human CMEC were treated with 10 μM A$\beta_{40}$-Q22, alone or in combination with 10 or 30, μM of 4ITP. Western blot analysis of tight junction proteins occludin and claudin-5 were evaluated at 6 hours. At 6 hours claudin-5 was significantly increased in the presence of Q22 + 4ITP in addition to 4ITP alone when compared to the control. Data represents 3 individual experiments of 2 replicates per group and graphed as mean ± SEM. Statistical significance was evaluated by One-way ANOVA followed by Tukey post-hoc test. *P< 0.05, **P< 0.01,****P<0.0001

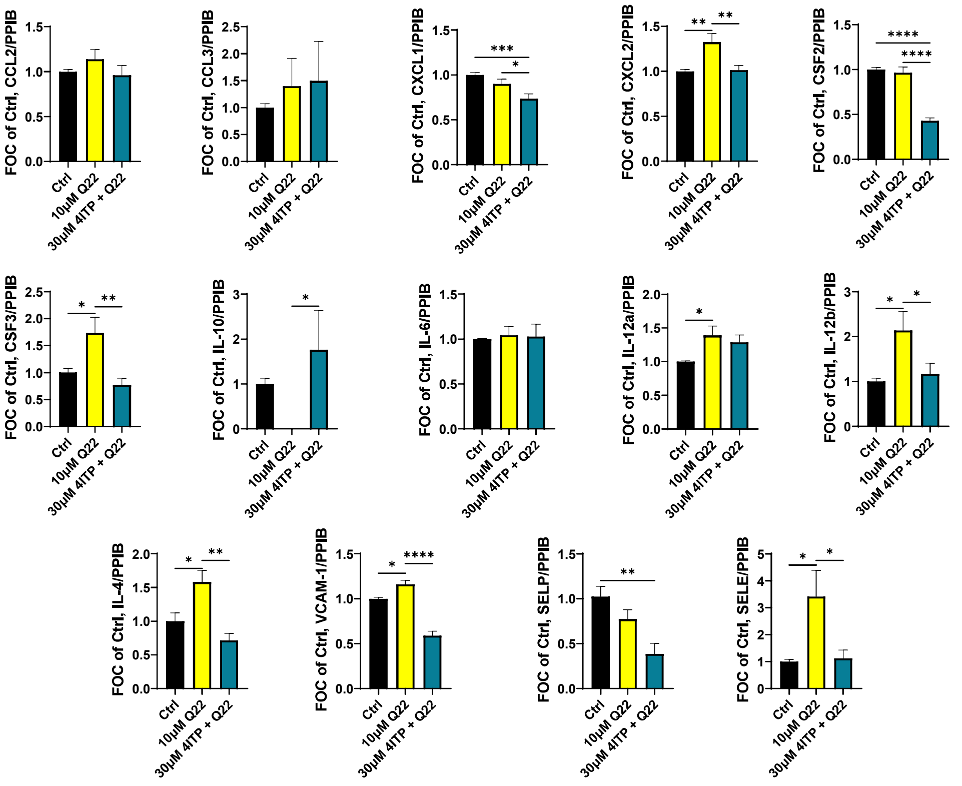


**Figure S3. CA-V inhibition prevents amyloid-induced endothelial cell activation in human CMEC**. Human CMEC were treated with 10 µM $A\beta_{40}$-Q22, alone or in combination with 30µM of 4ITP for 24h. The mRNA expression of 14 cytokines and chemokines was measured. The mRNA expression of CXCL2, CSF3, IL-12a/b, IL-4, VCAM-1 and E-selectin were all upregulated in the presence of amyloid. Co-treatment with 4ITP prevented the significant increase in CXCL2, CSF3, IL-12a/b, IL-4, VCAM-1, and E-selectin. Additionally, 4ITP significantly enhanced mRNA expression of IL-10 in the presence of amyloid. Finally, 4IP significantly reduced CSF2, and P-selectin. Data represents 3-4 individual experiments of 2 replicates per group and graphed as mean ± SEM. Statistical significance was evaluated by One-way ANOVA followed by Dunnet's post-hoc test. *P< 0.05, **P< 0.01,****P<0.0001

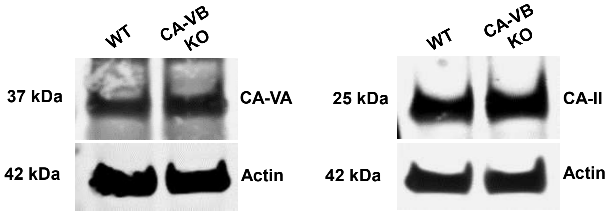


**Figure S4. Expression of CA-VA and CA-II isoforms in hCMEC/D3 CA-VB KO cells.** WB analysis confirmed CA-VA and CA-II normalized to loading control actin are expressed in hCMEC CA-VB KO cells.

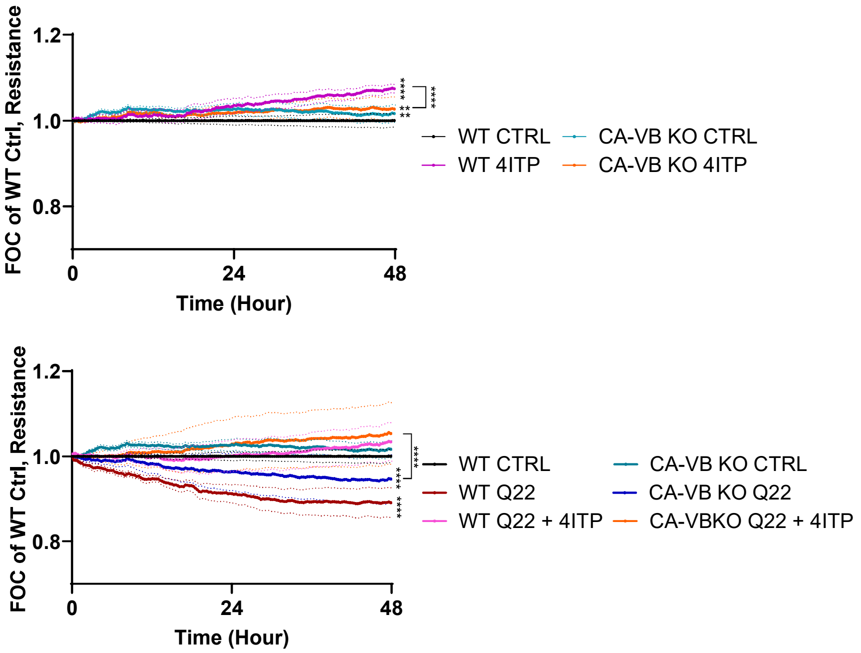


**Figure S5. 4ITP prevents Aβ40-Q22 induced loss in endothelial barrier integrity in CA-VB KO cells.** hCMEC barrier resistance was assessed by ECIS Zθ to evaluate barrier integrity. Human CMEC were treated with 10μM Aβ40-Q22 (Q22), alone or in combination with 30μM 4ITP. Data represents 3 individual experiments with 2 replicates per group, graphed as mean ± SEM. Statistical significance was evaluated by One-way ANOVA followed by Tukey post-hoc test. *P< 0.05, **P<0.01, ***P<0.001, ****P<0.0001

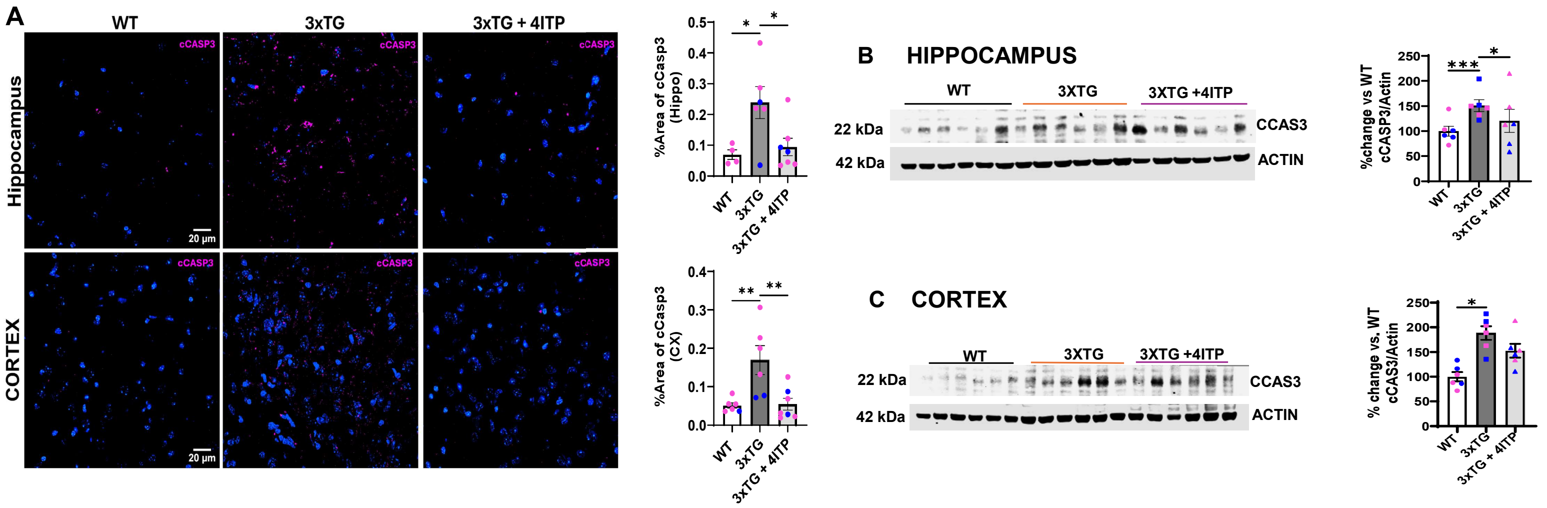


**Figure S6. 4ITP prevents caspase-3 activation in the brain of 3xTG mice.**. **(A)** Representative images of immunohistochemistry caspase-3 staining in the hippocampus and cortex of WT and 3xTG mice. Quantification to the right. Active caspase-3 is significantly increased in the hippocampus and cortex, and then significantly reduced with 4ITP treatment. **(B-C)** Western blot analysis of caspase-3 cleavage in the hippocampus and cortex with quantification to the right. ± SEM. Dots are pink and blue to represent female and female mice, respectively. WT N=4, 3xTG N= 6, 3xTG + 4ITP, N=7. Data represents 5-10 images per mouse per area, graphed as mean ± SEM. Statistical significance was evaluated by One-way ANOVA followed by Dunnett's multiple comparisons test. *P< 0.05, **P<0.01, ***P<0.001, ****P<0.0001

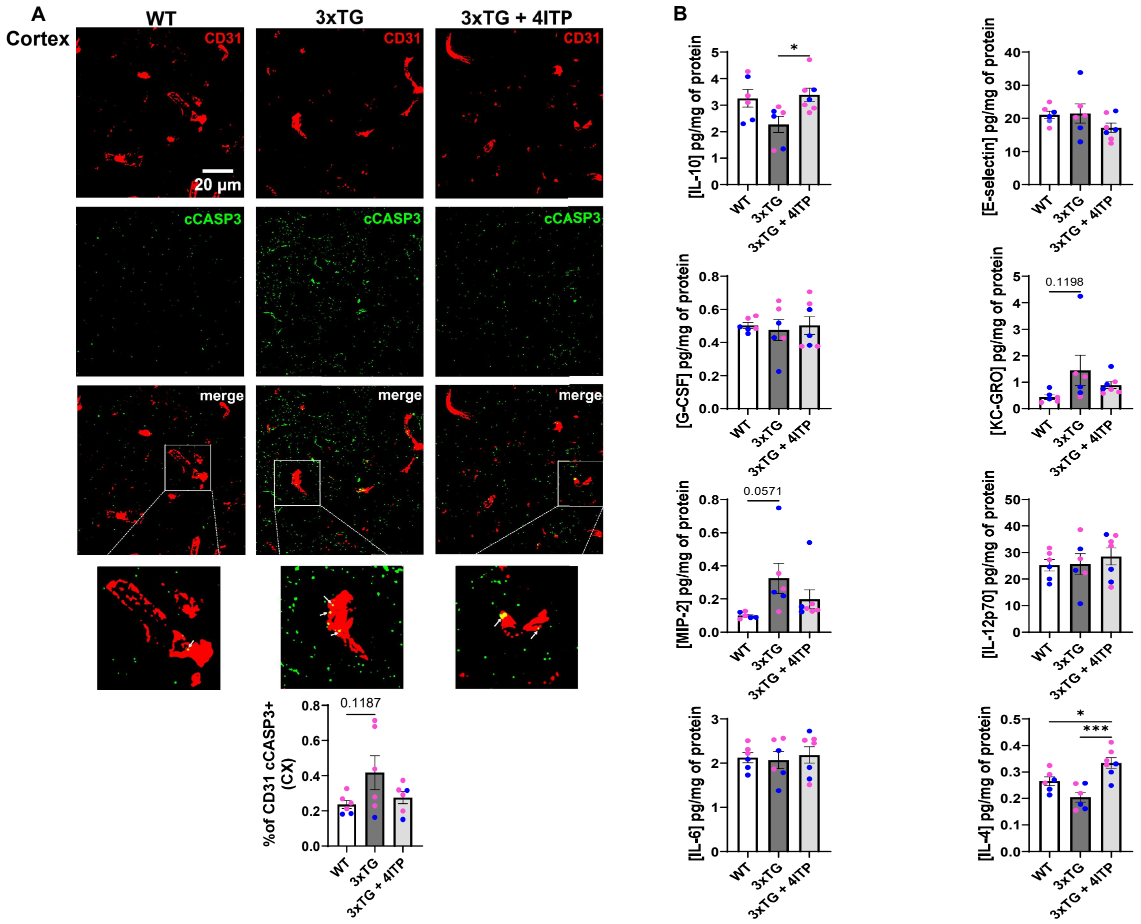


**Figure S7. 4ITP prevents endothelial cell apoptosis and mediates neuroinflammation in the cortex of 3xTG mice. (A)** Colocalization of CD31 and active caspase-3 was evaluated through immunohistochemistry. CD31 in red and cleaved caspase-3 in green. Colocalization is observed in yellow in the merged images. Quantification below panel. **(B)** Quantification of chemokines and cytokines in the cortex of WT, 3xTG and 3xTG treated mice through MSD . The reduction in IL-10 observed in 3xTG non-treated mice is significantly increased with 4ITP treatment, in addition, MIP-2 increase is mitigated with 4ITP. IL-4 reduction is significantly inhibited by 4ITP in 3xTG-treated mice. Pink and blue dots indicate female and male mice, respectively. (A) Data was analyzed as 5-10 images per mouse, graphed as mean ± SEM. (B) Data represents two replicates/ mouse graphed as ± SEM. Statistical significance was evaluated by One-way ANOVA followed by Tukey post-hoc test. *P< 0.05, **P<0.01, ***P<0.001, ****P<0.0001

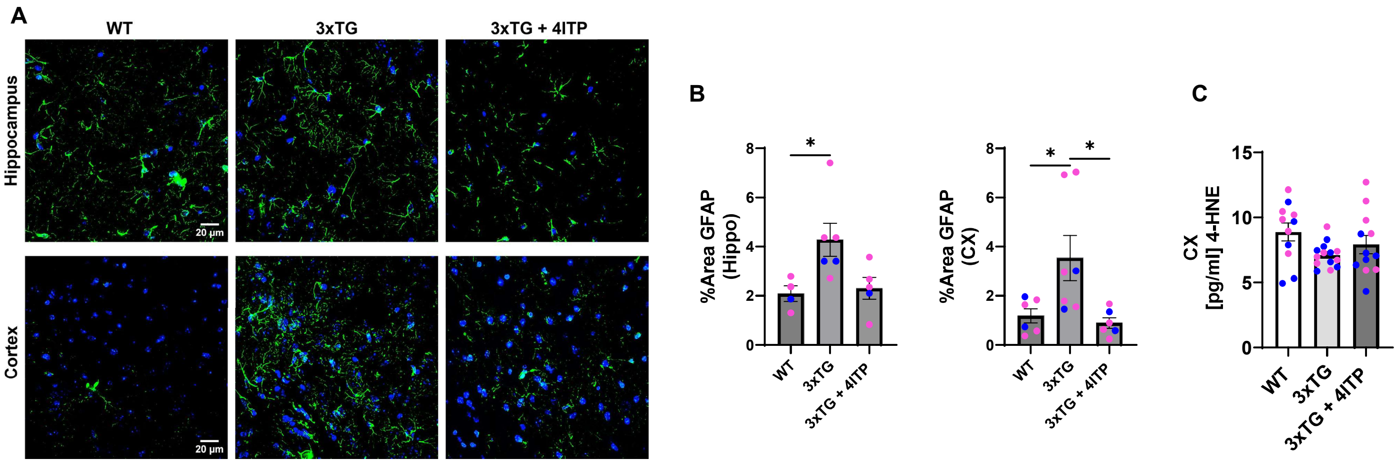


**Figure S8. 4ITP prevents astrocyte activation in hippocampus and cortex while 4-HNE levels stay the same in 3xTG mice. (A-B)** The levels of GFAP were evaluated through immunohistochemistry in AD pathological brain areas (hippocampus and cortex). We observed that the enhanced %area of GFAP is reduced with 4ITP treatment. This rescue is observed in both the hippocampus and cortex. WT N=4-6, 3xTG N= 6-7, 3xTG + 4ITP, N=5-6. Blue and pink symbols indicate male and female mice, respectively. Data represents 5-10 images per mouse per area, graphed as mean ± SEM. Statistical significance was evaluated by One-way ANOVA followed by Dunnett's multiple comparisons test. *P< 0.05 **(C)** 4-HNE was measured with an ELISA kit in the cortex of WT, 3xTG non-treated and treated mice (N=11-14/group) 4-HNE is not significantly changed. Male and female mice depicted with blue and pink dots respectively. Data represents 2 replicates per mouse, graphed as mean ± SEM. Statistical significance was evaluated by One-way ANOVA followed by Tukey post-hoc test.

.

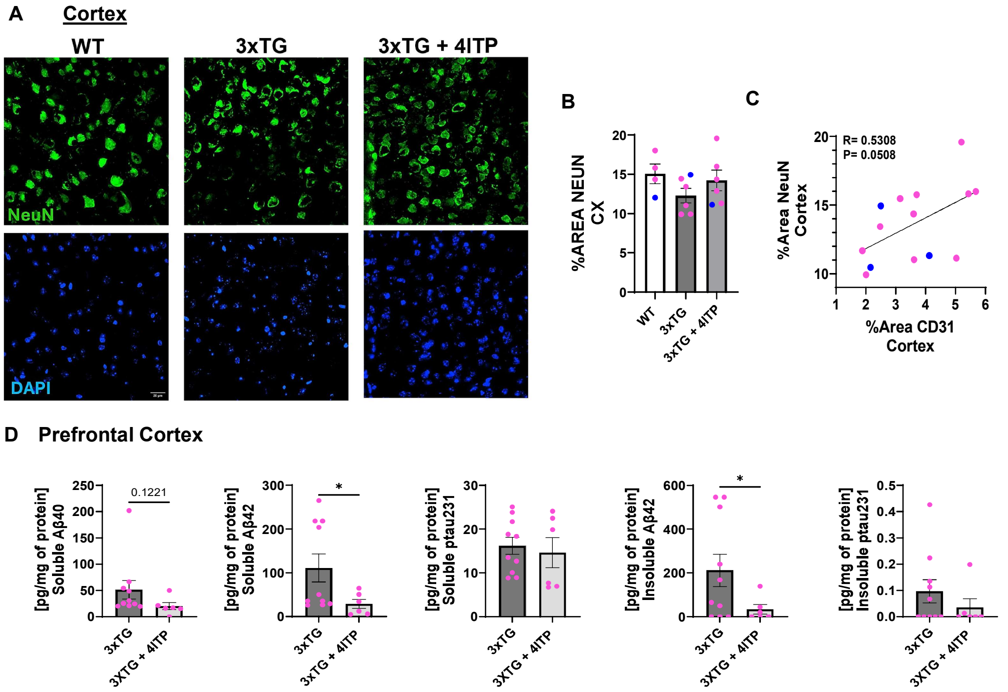


**Figure S9. 4ITP increases neuronal cells area, correlating with area of CD31+ cells. 4ITP decreases amyloid accumulation in the cortex of treated 3xTG mice. (A)** Representative IHC images of neurons marked with green and DAPI in blue in the cortex of WT and 3xtg mice. **(B)** 3xTG mice have a non-significant trend in decrease of %Area NeuN which appears improved with 4ITP. **(C)** Correlation between %Area NeuN and %Area CD31 is positive in the cortex of WT, 3xTG treated and non-treated mice (R=0.53, p= 0.05). Mice are indicated with pink and blue dots as female and male, respectively. **(D)** Soluble and insoluble amyloid and phospho-tau 231 quantification with ELISA in the prefrontal cortex of female 3xTG treated and non treated mice. 4ITP significantly reduced soluble and insoluble Abeta42 while Abeta40 was also trending in towards a reduction. (A-C) N=5-6 mice/group. Data was analyzed in 5-10 images per mouse, graphed as mean ± SEM. (D) Data represents two replicates/ mouse graphed as ± SEM. Statistical significance was evaluated by One-way ANOVA followed by Tukey post-hoc test. *P< 0.05, **P<0.01, ***P<0.001, ****P<0.0001

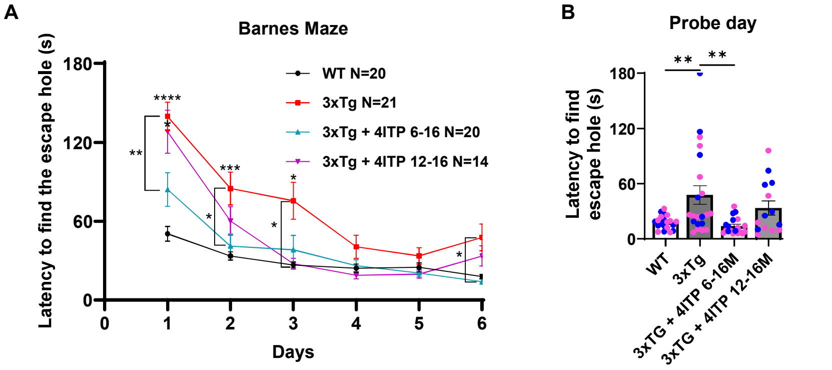


**Figure S10. Long treatment prevents cognitive decline more effectively than short treatment in 3xTG mice. (A**) Barnes maze at 15-16 months of age. Latency to find the escape hole was recorded in two sessions/day and averaged. On day 1 of the maze, the long treatment had significantly improved latency compared to 3xTG mice. This difference is observed throughout the training days, particularly at day 1-2 and 6. **(B)** On probe day, the increase in the latency to find the escape hole in 3xTG mice is significantly reduced to WT levels with the long treatment, while there is no significant difference between the 3xTG and short treatment. Male and female mice are indicated with blue and pink dots respectively. Statistical significance was evaluated by One-way ANOVA followed by Tukey multiple comparisons test. *P< 0.05. **P<0.01 ***P<0.001****P<0.0001

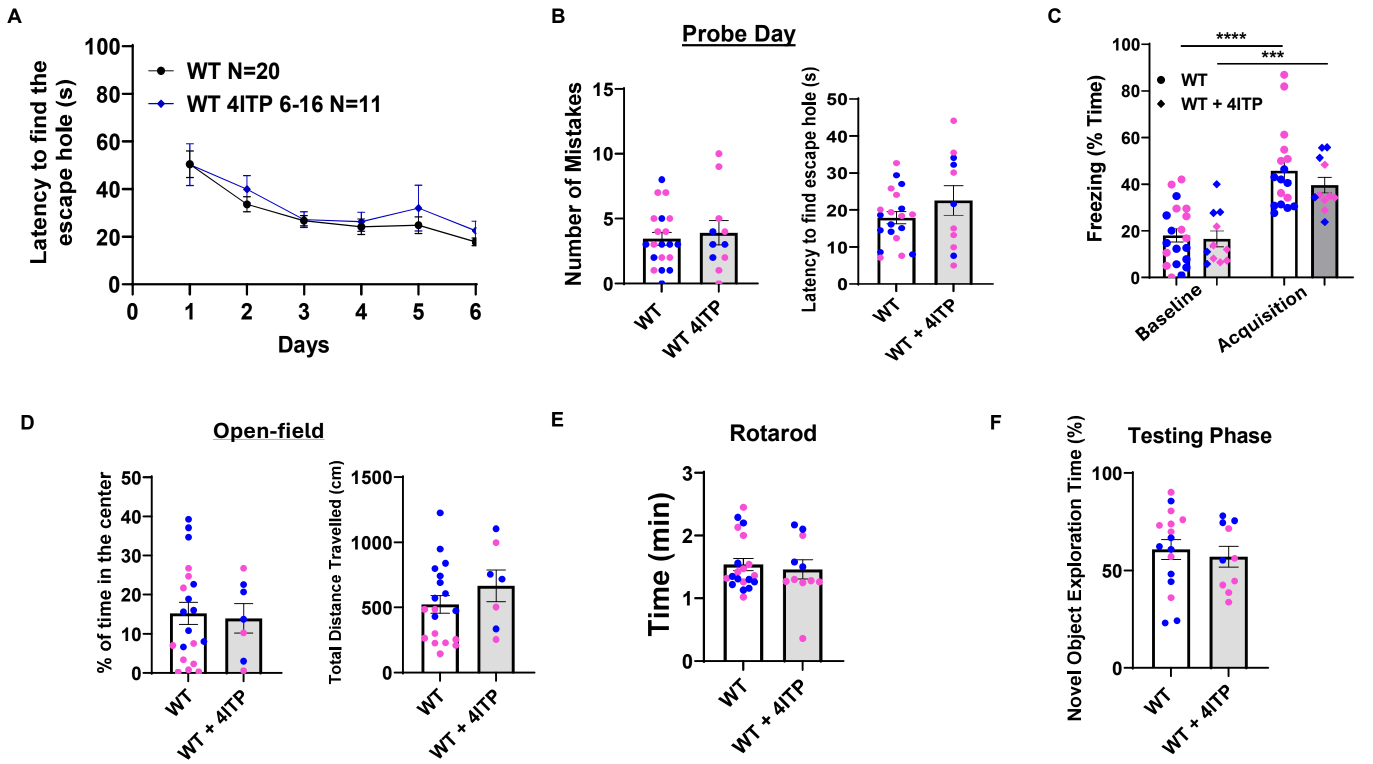


**Figure S11. Long-term treatment with 4ITP in WT mice did not alter spatial memory or locomotor activity. (A**) Barnes maze (WT, N=20 WT+4ITP N=7-11, males and females 15-16 months of age). Latency to find the escape hole was recorded two sessions/ day and averaged and then graphed per day. **(B)** There are no significant differences between treated and non-treated WT mice in the number of mistakes or the latency to find the escape hole on probe day. **(C)** There is a significant increase in freezing time in both treated and non-treated WT mice when comparing acquisition and baseline. **(D)** Motor function was assessed with open-field and **(E)** Rotarod. (**F**) There were no significant differences between WT treated and non-treated mice in the %of novel object exploration during the testing phase. **(B-F)** Male and female indicated with blue and pink dots respectively. Statistical significance was evaluated using Welch's T test. ***P<0.001

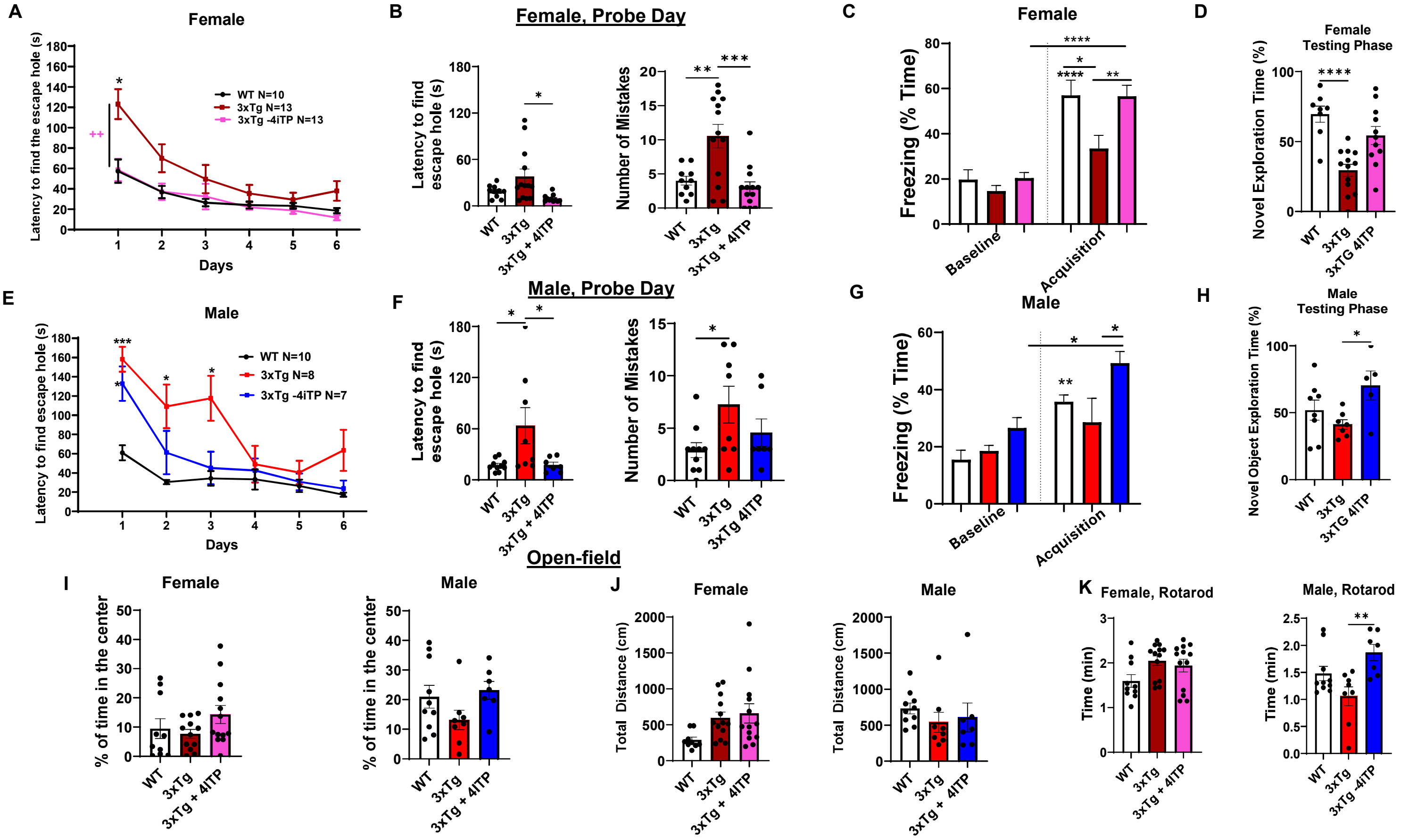


**Figure S12. Long-term treatment with 4ITP in 3xTG separated by sex in cognitive test, Barnes Maze, Fear-condition, and novel object recognition test. Additionally, locomotor tests rotarod and open-field**. **(A and D)** Barnes maze (females and males 15-16 months of age) Latency to find the escape hole was recorded two sessions/ day and averaged and then graphed per day. Female and male treated mice had improved spatial memory compared to non-treated 3xTG. **(B and F)** represent probe day latency to find the escape hole and number of mistakes. **(C and G)** Fear conditioning is rescued in both female and male mice represented through %freezing time during acquisition. **(D and H)** Novel object recognition test, Testing phase, indicates a mild rescue in females and males, significantly in males**. (I and J)** Open-field test demonstrates no motor impairments between groups, in addition to **(K)** Rotarod motor test in female and male mice. No significant changes in female while 4ITP improved motor function in male mice. Statistical significance was evaluated by One-way ANOVA followed by Tukey multiple comparisons test. *P< 0.05. **P<0.01 ***P<0.001
